# Global Profiling of 2-hydroxyisobutyrylome in Common Wheat

**DOI:** 10.1101/2020.03.05.978155

**Authors:** Ning Zhang, Lingran Zhang, Linjie Li, Junyou Geng, Lei Zhao, Yan Ren, Zhongdong Dong, Feng Chen

## Abstract

As a novel post-translational modification (PTM), lysine 2-hydroxyisobutyrylation (Khib) has been found to play a role in active gene transcription in mammalian cells and yeast, but the function of Khib proteins in plants remains unknown. In this study, we used western blot to demonstrate that Khib is an evolutionarily-conserved PTM in wheat and its donators, with the highest Khib abundance occurring in hexaploidy wheat. Additionally, global profiling using affinity purification and mass spectroscopy of 2-hydroxyisobutyrylome revealed that there were 3348 lysine modification sites from 1074 proteins in common wheat (*Triticum aestivum* L.). Moreover, bioinformatic data indicated that Khib proteins participate in a wide variety of biological and metabolic pathways. Immunoprecipitation and western blot confirmed that Khib proteins had an *in vivo* origin. A comparison of Khib and other major PTMs revealed that Khib proteins were simultaneously modified by multiple PTMs. Using mutagenesis experiments and Co-IP, we demonstrated that Khib on K206 is a key regulatory modification of phosphoglycerate kinase enzymatic activity and found that de-Khib on K206 affects protein interactions. Furthermore, Khib production of low-molecular-weight proteins was a response to the deacetylase inhibitors nicotinamide and trichostatin A. This study provides evidence that enhances our current understanding of Khib in wheat plants, including the cooperation between this PTM and metabolic regulation.

## Introduction

Protein post-translational modification (PTM) can change the charge, conformation, and molecular weight of proteins by adding chemical groups to the amino acid residues of a protein that can also expand the biological functions of the protein [1]. PTMs have been found to play a vital role in diverse biological processes by regulating protein function [2]. Lysine acetylation (Kac) is an important PTM that neutralizes positively-charged lysine residues; previous studies of Kac have mainly focused on nuclear proteins [3, 4]. Using mass spectrometry, a high abundance of non-histone proteins has been extensively characterized recently [5–9]. Kac is currently known to regulate diverse protein properties, including subcellular localization, DNA-protein interactions, protein stability, protein-protein interactions, and enzymatic activity [7, 8, 10, 11]. With the help of high sensitivity mass spectrometry, there are currently 9 novel lysine PTMs reported, including formylation (Kfor), crotonylation (Kcr), butyrylation (Kbu), succinylation (Ksu), malonylation (Kma), propionylation (Kpr), glutarylation (Kglu), *β*-hydroxybutyrylation (Kbhb), and 2-hydroxyisobutyrylation (Khib), and the catalog is still growing [12–18]. These novel PTMs have been mainly found in mammalian and yeast cells, and also in plants, such as *Arabidopsis thaliana* [19], rice [20–23], and tobacco [19, 24]. Increasing evidence suggests that these new types of PTMs are involved in multiple kinds of cellular and metabolic pathways in plants [20].

The wheat genome is large and advancements have been slower compared with other plants, but some progress has been made. To date, only two novel PTMs, Ksu and Kma, have been identified in wheat [25–27]. These new acylations were initially identified in the ε-amino group on the lysines of core histones [25, 26], while recent studies have indicated nuclear, cytoplasmic, and mitochondrial proteins can also harbor lysine acylations [28]; this is because all acylation types require acyl-CoAs as corresponding donors. In addition, lysine PTMs have been found to participate in diverse cellular metabolic pathways [2, 18, 29–34]. Altogether, these studies have dramatically enriched our understanding of histone modifications. As a next step, many researchers have started investigating the potential function of histone modifications in transcription as well as the enzymes involved in adding (writers), removing (erasers), and reading (readers) PTMs.

As a new type of histone marker, Khib was found to be conserved in the genomes of many species including yeast and humans. At present, Khib has been initially identified in 63 lysine sites on human and mouse histones and could play a critical role in the regulation of chromatin functions [16]. Khib is also important for histone 4 lysine 8 (H4K8), as H4K8hib is associated with active gene transcription in meiotic and post-meiotic cells [16]. Moreover, a recent study indicated that Khib on H4K8 is regulated by glucose homeostasis in *Saccharomyces cerevisiae*, where histone lysine deacetylases Rpd3p and Hos3p function as regulatory enzymes for lysine de-Khib reactions for H4K8 [35]. Eliminating Khib at this site resulted in a reduced life span, implying that the modification of the H4K8 site may be involved in aging [35].

Over the past several years, many regulatory enzymes (writers and erasers) have been identified [36]. A recent study indicated that histone deacetylase 2 (HDAC2) and histone deacetylase 3 (HDAC3) were acting as “erasers” in mammalian cells, and Esa1p and its homologue Tip60 were discovered to be “writers” in budding yeast and in human cells, respectively [37]. Another recent study identified the histone lysine acetyltransferase (HAT) EP300 was a ‘‘writer’’ for Khib in mammalian cells, and that EP300 was able to regulate glycolysis through Khib of glycolytic enzymes, as well as mediate cell survival via nutritional regulation through glycolysis [37]. A very recent study indicated that CobB serves as a de-Khib enzyme that regulates glycolysis and cell growth in bacteria [38]. Proteome-wide profiling of the Khib in mammalian cells, yeast, *Proteus mirabilis*, rice, and *Physcomitrella patens* has discovered several nuclear, cytosolic, and mitochondrial proteins with acetyllysine modifications, which provided evidence that Khib may play a vital role in cell metabolism; this has stimulated research on the non-nuclear functions of the PTM [23, 35, 39–41]. Although there has been progress, it has been mainly in mammalian cells and yeast, while studies in plants are still limited. Therefore, it is interesting to determine the function of Khib proteins in plants.

In this study, we analyzed wheat Khib sites by utilizing high performance liquid chromatography (HPLC) and tandem mass spectroscopy (LC-MS/MS). A total of 3348 lysine modification sites on 1074 proteins were identified in common wheat. Using mutagenesis experiments, we confirmed that Khib on K206 of phosphoglycerate kinase (PGK) was a key regulatory modification; we also observed PGK protein interactions impacted by Khib. Western blot results indicated that wheat sirtuin or Zn-binding HDAC proteins regulated the Khib of low-molecular-weight proteins. Our data indicated that the broad regulatory scope of Khib was comparable with that of other major PTMs. These novel Khib sites and proteins not only expand our understanding of the functional roles of Khib but also lay a foundation for future studies in plants.

## Results and discussion

### Evolutionary conservation of Khib

Previous studies have indicated that histone Khib is an evolutionarily-conserved PTM in eukaryotic cells (e.g., HeLa cells), including Drosophila S2, mouse embryonic fibroblasts, and yeast *Saccharomyces cerevisiae* cells [16]. To test whether Khib is present in wheat and its donor progenitors, the pan-anti-2-hydroxyisobutyryllysine (pan-anti-Khib) antibody was used for western blot analyses in *Triticum urartu* (PI428185, AA), *Aegilops speltoides* (PI542245, SS), *Aegilops tauschii var. strangulate* (AS2393, DD), tetraploid *Triticum durum* Langdon (AABB), and hexaploidy wheat AK58 (AABBDD) (**Figure 1**). Our analysis revealed that Khib was widely distributed and mainly enriched in 40–70 kDa in hexaploidy wheat and its progenitors. Hexaploidy wheat had the highest Khib levels compared with its progenitors. Additionally, the Khib level of the diploid progenitors (AA, SS, and DD) was lower than the tetraploid progenitor (AABB). These results indicate that Khib is an evolutionarily-conserved PTM in wheat and its donators.

**Figure 1.**
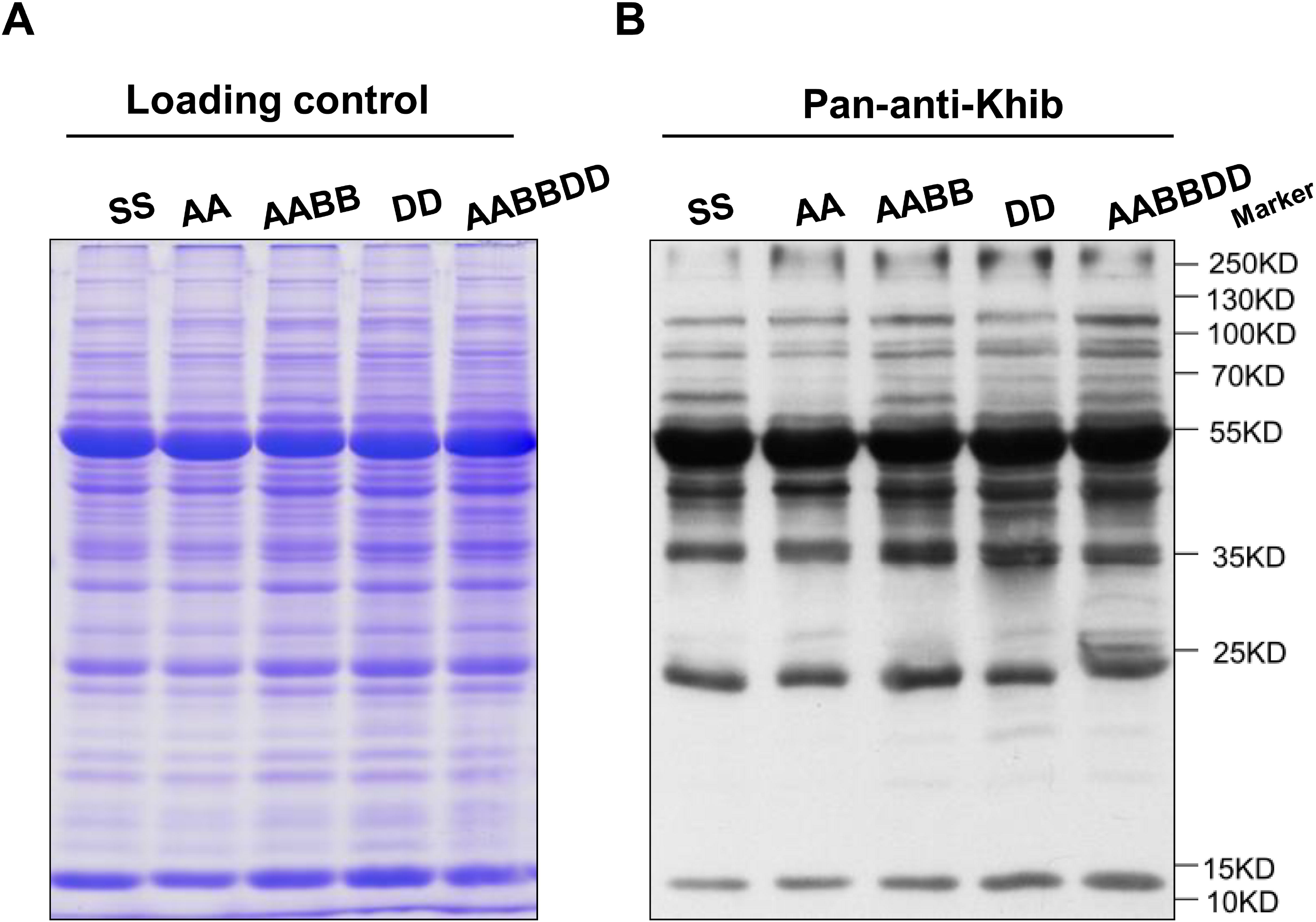
Khib is an evolutionarily-conserved post-translational modification. (A) SDS-PAGE gel. (B) Western blot of the Khib proteins in wheat and its donor progenitors. SS: *Aegilops speltoides*; AA: *Triticum Urartu*; AABB: *Triticum durum*; DD: *Aegilops tauschii*; AABBDD: *Triticum aestivum* L. Khib, lysine 2-hydroxyisobutyrylation; Pan-anti-Khib, pan-anti-2-hydroxyisobutyryllysine antibody. The same below.

### Global analysis of 2-hydroxyisobutyrylome in common wheat

#### The lysine 2-hydroxyisobutyrylome profile

A proteome analysis was conducted using the antibody-based affinity enrichment and LC-MS/MS methods. A total of 3348 Khib sites of 1074 proteins in hexaploidy wheat were identified (**Figure 2A**; Supplemental Table S1, and Figure S1). Most of the mass errors were less than 5 ppm, confirming the high accuracy of our MS data (Supplemental Figure S2). The length of most peptides (98%) was in line with the properties of tryptic peptides (from 7 to 28 amino acids) (**Figure 2B**). To evaluate the coverage of Khib in substrate proteins, we counted the number of modification sites per protein. Of the 1074 Khib proteins, 45.3% and 43.1% have 1 and 2–6 Khib sites, respectively, while the remaining 11.6% have ≥7 Khib sites (**Figure 2C**), indicating that 54.7% contained multiple Khib sites. Wheat 2-hydroxyisobutyrylome was larger than what has been previously reported for acetylome, malonylation, succinylome, and ubiquitome in plants [25–27, 42]. This is in agreement with a previous study, which indicated there are more Khib sites on histones than on other PTM sites [16], implying that lysine 2-hydroxyisobutyrylaton was an abundant PTM and may play a vital role in substrate protein regulations.

**Figure 2.**
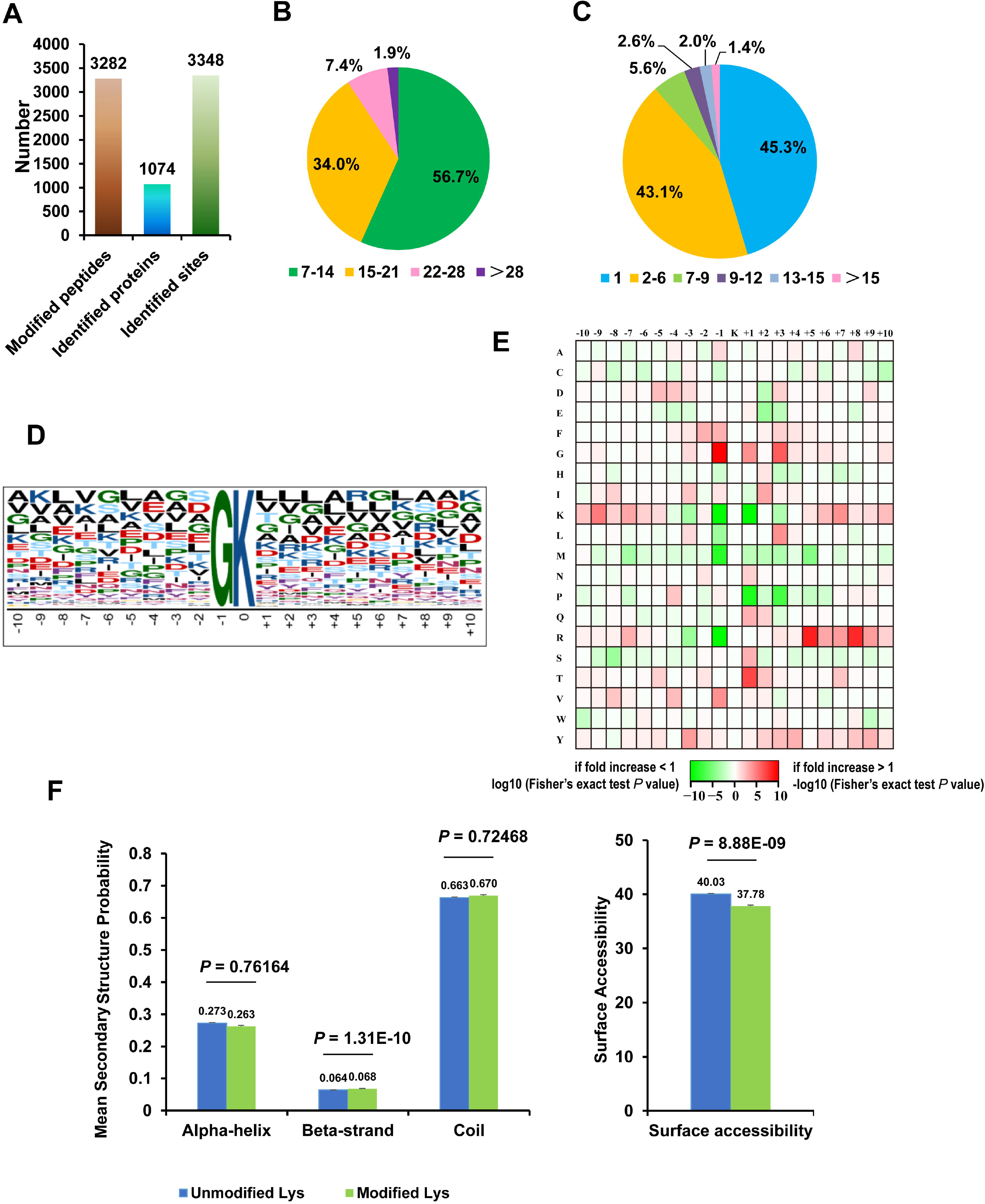
Proteome-wide identification and properties of Khib sites in wheat. (A) Statistical analysis of the Khib sites, proteins, and peptides. (B) Peptide length distribution and percentage. (C) The number and proportions of Khib sites per protein. (D) Khib motifs and the conservation of Khib sites. The height of each letter represents its frequency, and the central K represents the Khib lysine. (E) Heat map of amino acid compositions of the Khib sites. (F) Probabilities of Khib in three different protein secondary structures (alpha-helix, beta-strand, and coli; left) and the predicted surface accessibility of Khib sites (right). Lys: lysine.

Sequence motifs in Khib peptides were determined by flanking the sequence of these sites using the Motif-X program. According to the heat map of amino acid compositions at the Khib sites, considerable non-polar hydrophobic amino acid glycine (G) was enriched at the −1 position, indicating that there was likely Khib at this location in wheat (**Figure 2D**–**E**). This pattern was different from Khib-modified motifs in rice, *Proteus mirabilis*, and humans [23, 39, 40], although GKhib simultaneously exists in *Physcomitrella patens* proteins [41]. This surprising result indicates the complexity of underlying PTM mechanisms in plants.

We conducted a structural analysis of all identified proteins using NetSurfP. Results indicated that 26.3% and 6.8% of the Khib sites were located in the *α*-helix and *β*-strand, respectively, while 6.7% of the Khib sites were located in disordered coils (Supplemental Table S2; **Figure 2F**). Khib sites were found more frequently in the *β*-strand regions (*P* = 1.31e-10) and less frequently in the *α*-helix (*P* = 0.762) and disordered coil (*P* = 0.726) regions when compared with unmodified lysine residues. Thus, it is clear that Khib has a preference for secondary structures. Identified Khib sites were further evaluated for solvent accessibility, and it was found that 37.8% of the Khib sites were exposed to the protein surface, compared with 40.0% of those on unmodified lysine residues (*P* = 8.88e-09) (Supplemental Table S2; Figure 2F). In comparison with unmodified counterparts, lysine residues at Khib sites were less surface-accessible. The lower surface accessibility of the Khib sites implies that lysine Khib may occur in a selective process.

#### Enrichment analysis of the Khib proteins

To evaluate Khib proteins, we performed the gene ontology (GO) functional classification according to their subcellular location, cellular component, molecular function, and biological processes (Supplemental Tables S3–S4 and Figure S3A–C). Based on subcellular location, there were 49.2%, 26.4%, 9.9%, and 6.4% Khib proteins in the chloroplast, cytoplasm, nucleus, and mitochondria, respectively, indicating a wide distribution of Khib proteins in wheat. Based on cellular component, Khib proteins were distributed in the cells (35.4%), the organelles (24.5%), the macromolecular complexes (21.8%), and membranes (17.3%). Based on molecular function analysis, 41.8% and 38.8% of Khib proteins were associated with binding and catalytic activities, respectively; this could imply that Khib is an important PTM in DNA transcription or protein interactions and has a large influence on metabolic processes. Biological process analysis showed that 36.8%, 30.7%, 19.6%, and 4.8% of Khib proteins were involved in metabolism, cellular processes, single organism proteins, and location, respectively. These features are similar to the previous reports in rice and *Physcomitrella patens*.

To further understand the characteristics and potential roles of wheat Khib proteins, we performed analyses of the enrichment of the GO term, Kyoto Encyclopedia of Genes and Genomes (KEGG) pathway, and protein domain (Supplemental Tables S5–S8 and **Figure 3A–C**). Biological process enrichment indicated that Khib proteins were enriched in a variety of cellular and metabolic processes, indicating a wide impact of this novel PTM in wheat. Similar to Kac, Khib of metabolically-related enzymes could have a conserved regulation pattern in different organisms [23, 35, 39–41]. KEGG enrichment analysis demonstrated that Khib proteins were enriched in multiple primary metabolic pathways (Figure 3B), such as carbon metabolism, carbon fixation in photosynthetic organisms, and photosynthesis-antenna proteins. In contrast, ribosomes, biosynthesis of amino acids, proteasomes, and arginine biosynthesis were also significantly enriched in the KEGG pathway analysis, suggesting the potential role of protein Khib in the regulation of protein biosynthesis and degradation. Protein domain enrichment analysis showed Khib proteins were enriched in the chlorophyll a/b binding protein domain, NAD(P)-binding domain, thioredoxin-like fold, as well as heat shock protein 70kD, histone H2A/H2B/H3, and pyridoxal phosphate-dependent transferase (Figure 3C). Enrichment analysis of KEGG pathways using the InterPro Domain protein database indicated that photosynthesis and carbon metabolism are likely significantly regulated by Khib.

**Figure 3.**
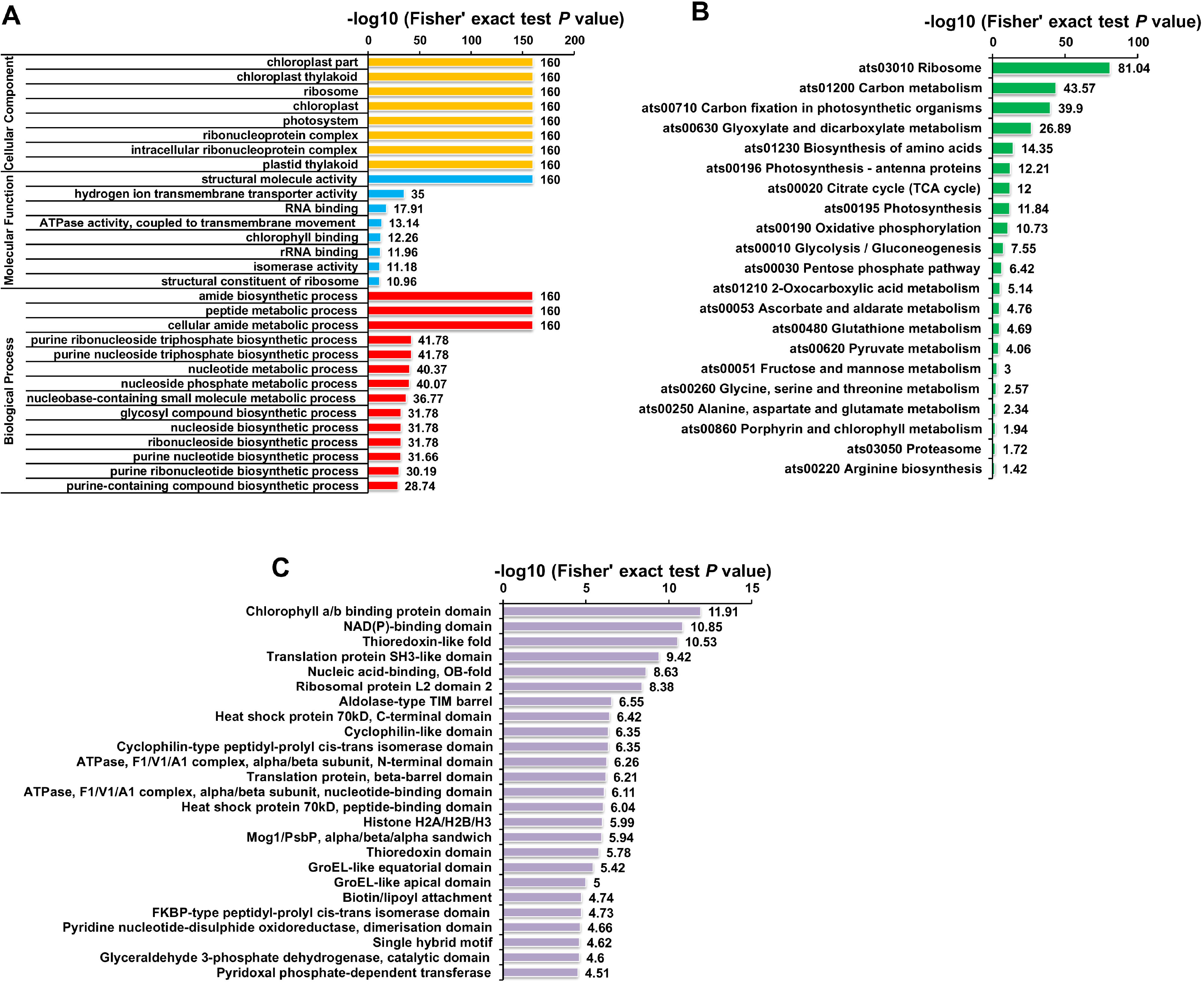
Enrichment analysis of Khib proteins in wheat. (A) GO analysis. (B) KEGG pathway. (C) Protein domain. GO, Gene Ontology; KEGG, Kyoto Encyclopedia of Genes and Genomes (the same below).

#### Interaction network of the Khib proteins

To quantify protein-interaction properties of the 2-hydroxyisobutyrylome, physical and functional interaction analysis was performed using STRING. We found the existence of connections among 435 Khib proteins, indicating Khib proteins are able to participate in a diverse set of functions in wheat (Supplemental Table S9 and Figure S4A–D). We also retrieved 21 highly interconnected clusters of Khib proteins using the Cytoscape software algorithm. For example, 102 ribosome-related proteins (Cluster I), 11 proteasome-related proteins (Cluster II), and 10 proteins associated with metabolic pathways (Cluster III) demonstrated the presence of close interaction networks (Supplemental Figure S4A–C). The pathways subnetworks suggested that all of 435 Khib proteins participated in a dense protein interaction network.

#### Immunoprecipitation (IP) and the Wheat Germ Protein Expression System confirmed the existence of Khib

Six proteins were identified, and their Khib states were independently confirmed by immunostaining (**Figure 4A**–**B**; Supplemental Table S1; **Table 1**). The large subunit of ribulose-1,5-bisphosphate carboxylase (RbcL) from wheat leaf proteins was immunoprecipitated by an anti-RbcL antibody, where Khib was clearly present (Figure 4A). Five proteins, i.e., catalase (CAT), 14-3-3, glutathione S-transferase (GST), PGK, and glyceraldehyde-3-phosphate dehydrogenase (GAPDH) from the wheat germ protein expression system (cell free) were expressed as fusions with the His-tag and/or Halo-tag and showed clear Khib signals; the positive control bovine serum albumin (BSA, Sigma) did not have a signal (Figure 4B). Together, these results demonstrate that 2-hydroxyisobutyrylome has an *in vivo* origin.

**Figure 4.**
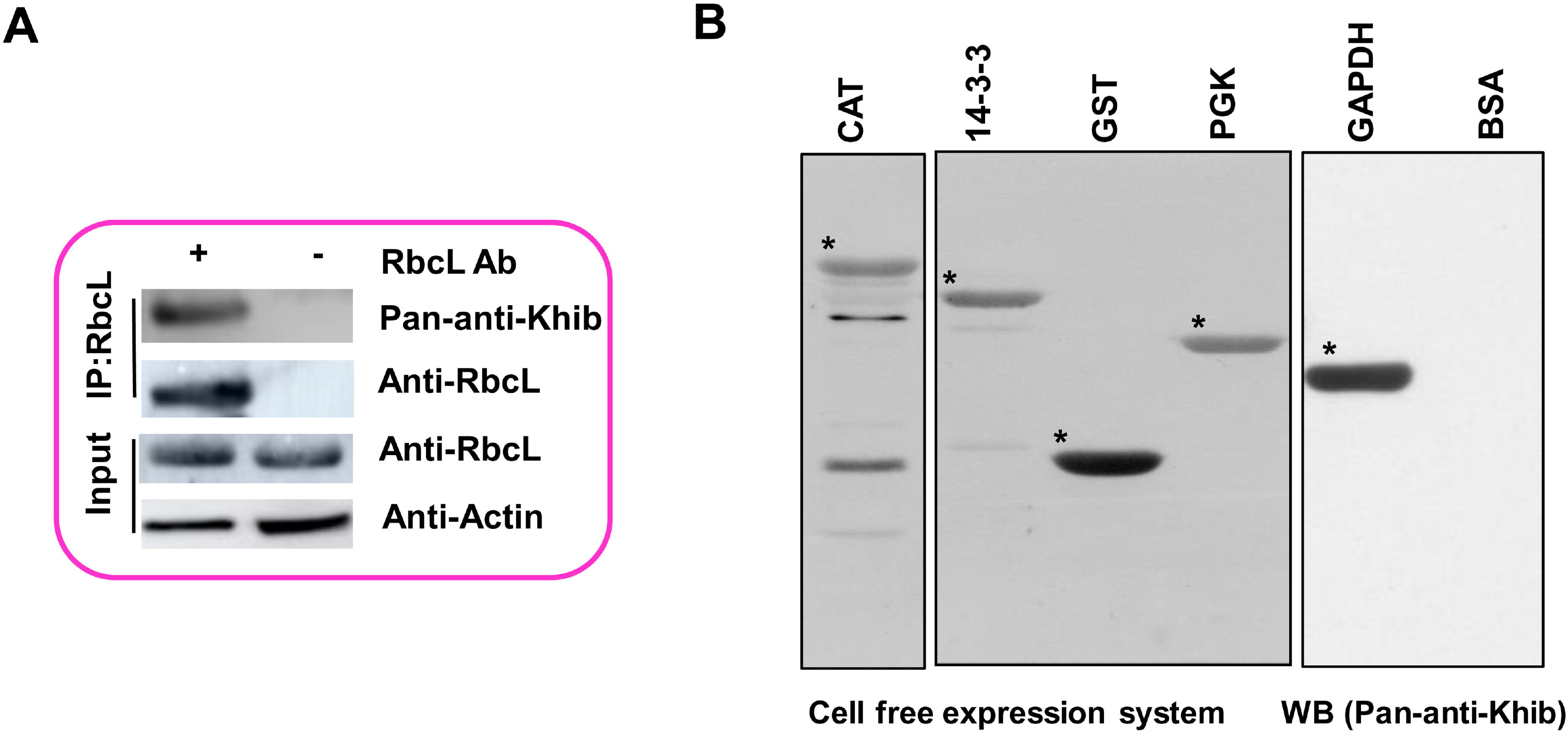
Independent validation of Khib of proteins. (A) IP of RbcL was performed with (+) or without (−) RbcL antibodies; eluted proteins were probed with either pan-anti-Khib or RbcL antibodies. (B) Purified recombinant protein and the positive control BSA were measured by western blot. IP, immunoprecipitation; RbcL, the large subunit of ribulose-1,5-bisphosphate carboxylase; CAT, catalase; GST, glutathione S-transferase; GAPDH, glyceraldehyde-3-phosphate dehydrogenase; PGK, phosphoglycerate kinase (the same below); BSA, bovine serum albumin (the same below).

**Table 1.** Khib sites information of proteins that used for IP and wheat germ protein expression system.

| Protein accession | Position | Amino acid | Protein description |
| --- | --- | --- | --- |
| P11383 | 7, 14, 18, 21, 32, 99, 146, 164, 175, 177, 183, 201, 227, 236, 252, 305, 316, 334, 356, 450 | K | Ribulose biphosphate carboxylase large chain |
| A0A1D5YMQ8 | 50, 72, 125, 163, 223, 227, 401, 424, 429, 464, 481 | K | Catalase |
| L0GED8 | 36, 56, 75, 89, 101, 124, 129, 142, 148, 250 | K | 14-3-3 protein |
| W5H4V7 | 35, 58, 78, 137, 202, 206, 366 | K | Phosphoglycerate kinase |
| A0A1D5TTT4 | 89, 96, 109, 121, 124, 159, 166, 167, 217, 222, 274, 276, 300, 301 | K | Glyceraldehyde-3-phosphate dehydrogenase |
| Q9SP56 | 138, 140, 152 | K | Glutathione S-transferase |

#### Conservation and uniqueness of the Khib proteins

To reveal the conservation and uniqueness of Khib among rice (*Oryza sativa*) [23], *Physcomitrella patens* [41], and wheat, we utilized a BLAST search to estimate the degree of conservation and uniqueness of the Khib proteins [42]. As shown in **Figure 5A**, 752 (70.0%) of the Khib proteins were orthologous proteins between wheat and the other two species (Supplemental Table S10; Figure 5A). A total of 178 Khib proteins were found in all three species, and KEGG enrichment analysis indicated that these orthologous proteins were abundantly enriched in ribosomes, carbon metabolism functions, the citrate cycle (TCA cycle), carbon fixation in photosynthetic organisms, and proteasomes (Supplemental Table S11; **Figure 5B**), suggesting the vital role and conservation of Khib in these different plant species. Among the 752 identified Khib proteins in wheat, 460 and 114 had conserved orthologs in *Physcomitrella patens* and rice, with an averaged similarity of 66.3% and 76.6%, respectively (Supplemental Table S10; Figure 5A). Moreover, 322 Khib proteins were unique in wheat and mainly enriched in proteins associated with carbon fixation in photosynthetic organisms, carbon metabolism, photosynthesis-antenna proteins, and photosynthesis (Supplemental Table S11; **Figure 5C**), indicating photosynthesis and carbon metabolism are likely significantly regulated by Khib in wheat.

**Figure 5.**
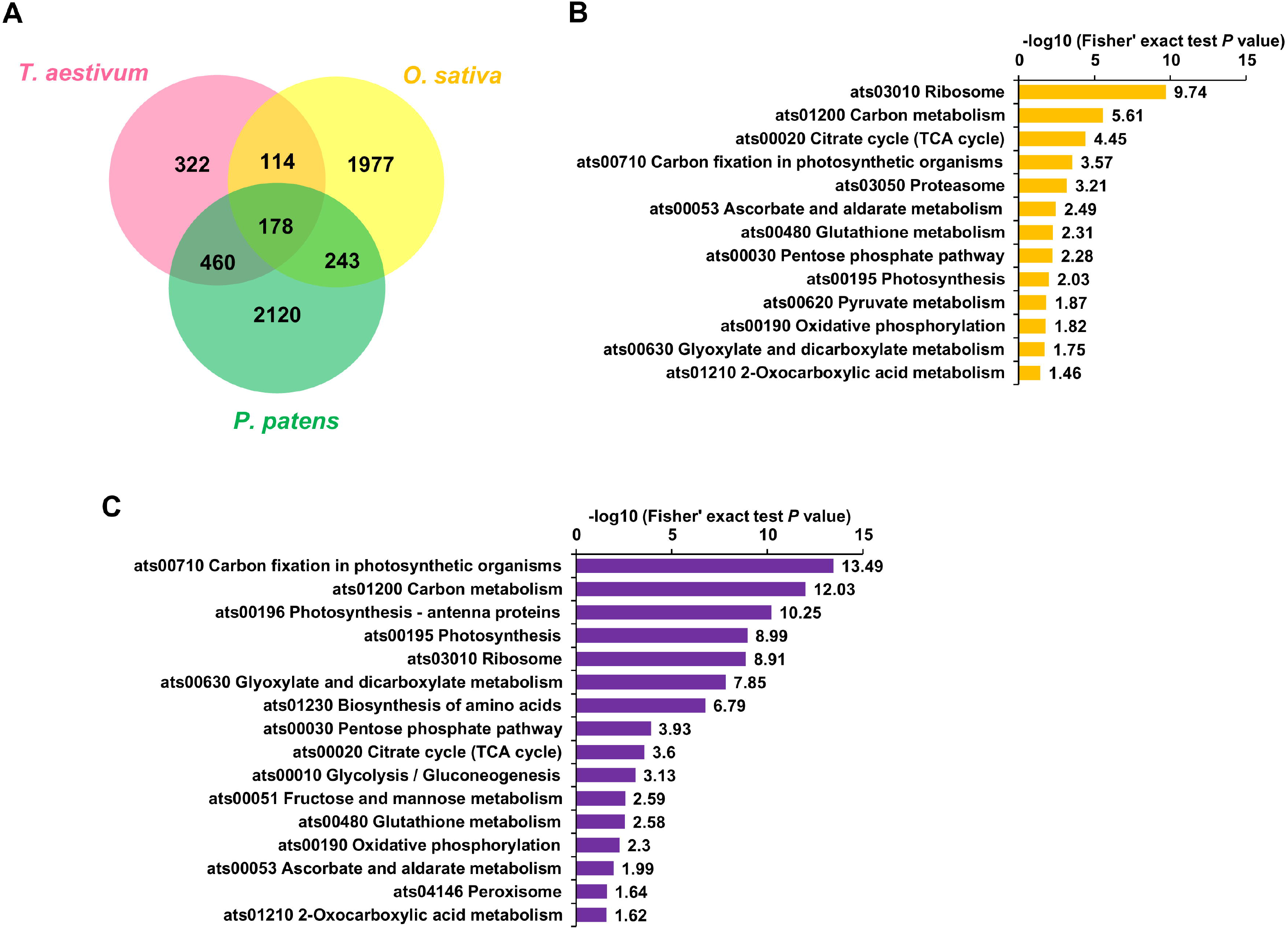
Conservation of Khib proteins in plants and uniqueness in wheat. (A) All identified Khib proteins in wheat compared with *Oryza sativa* and *Physcomitrella patens*. (B) KEGG enrichment of the common Khib proteins identified in wheat, *Oryza sativa*, and *Physcomitrella patens*. (C) KEGG enrichment of the uniqueness Khib proteins in wheat.

#### Overlap of Khib, Kma, Ksu, and Kac

Previous studies indicated that PTMs play vital roles in numerous biological processes and may have the same or distinct types of modifications [35, 43–45]. To determine whether the same lysine residues were modified by Khib and other PTMs simultaneously, we compared our Khib data with published Kma, Ksu, and Kac data in common wheat [25–27]. Results revealed that two proteins were modified by four PTMs. Namely, the 3, 9, and 5 Khib proteins were modified by Kma and Kac, Ksu and Kac, and Ksu and Kma, respectively. Additionally, the 12, 16, and 19 Khib proteins were modified by Kma, Ksu, and Kac, respectively (**Figure 6A**–**B**; Supplemental Tables S12–S13). In addition, the 19, 32, and 29 Khib sites were found at the same position as the Kma, Ksu, and Kac sites, respectively, while the 4, 3, and 5 Khib sites were found at the same position as the Kac and Kma, Ksu and Kma, and Ksu and Kac sites, respectively. Some of these Khib sites occurred at key function regions for proteins and shared the same lysis sites with other PTMs, which indicates that Khib likely possessed an important biological function.

**Figure 6.**
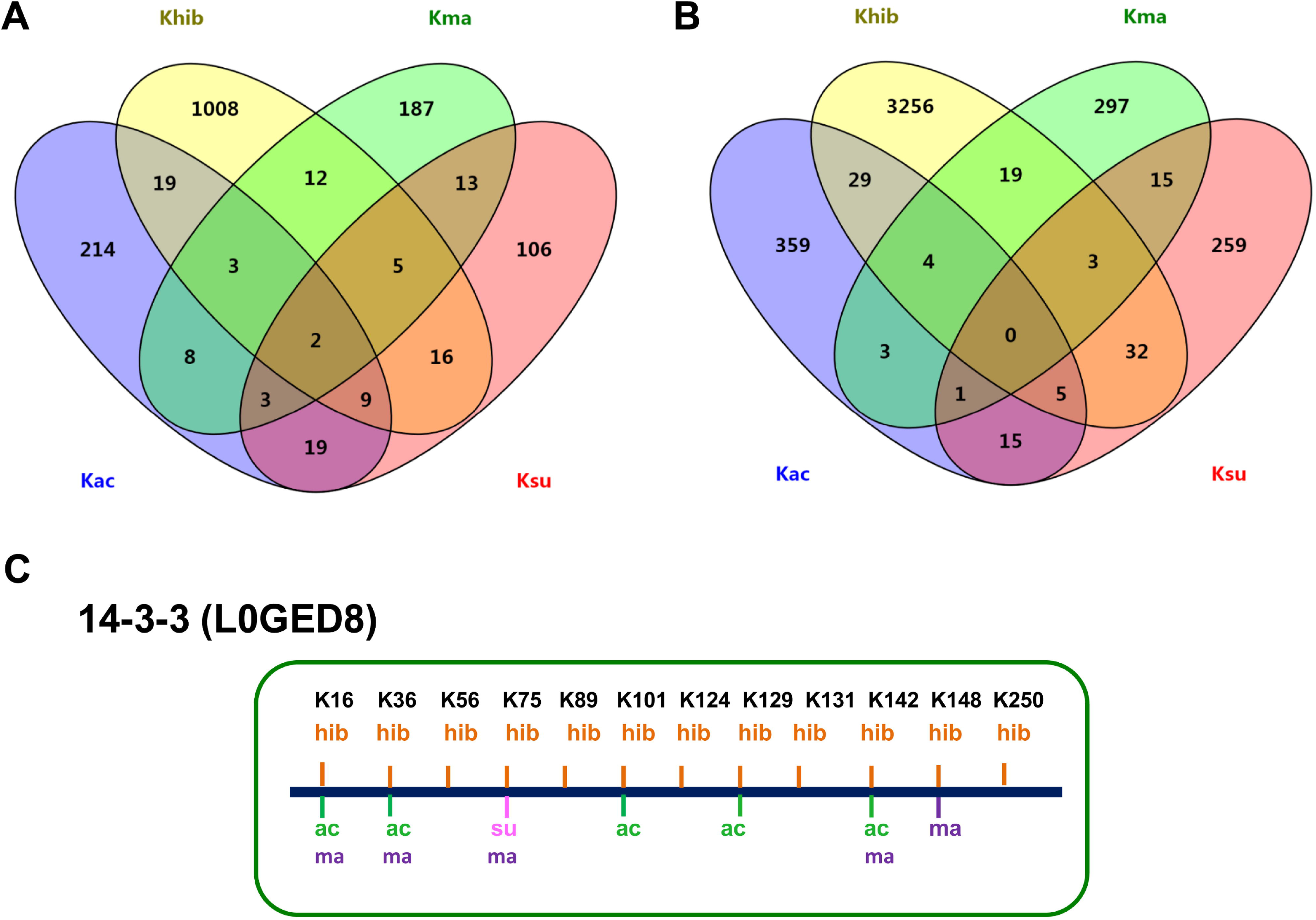
Overlap of wheat acetylome, succinylome, malonylome, and 2-hydroxyisobutyrylome. (A) Overlap of the number of proteins. (B) Overlap of the number of sites. (C) A representative protein 14-3-3 (L0GED8) showing the overlap of multiple post-translational modification sites. Ksu, lysine succinylation; Kma, lysine malonylation; Kac, lysine acetylation (the same below).

For example, in protein 14-3-3 (L0GED8), ten Khib sites were determined, and three sites at K16, K36, and K124 were found to be modified by Kac and Kma (**Figure 6C**). Specifically, one site at K75 was modified by Kma and Ksu simultaneously, K101 and K129 were modified by Kac, and K148 was modified by Kma. The tertiary structure of 14-3-3 was predicted by SWISS-MODEL, which confirmed the importance of K56 and K129 binding to their phosphopeptide ligands (Supplemental Figure S5). The Khib proteins associated with the Calvin-Benson cycle, transporter, glycolysis, and ribosome were also found to be Kac, Ksu, or Kma. These findings suggest that the same protein or lysine residue have multiple PTMs, and their combinatorial effects may regulate the function of wheat proteins.

#### Khib proteins involved in photosynthesis

Khib is known to occur on nuclear, cytoplasmic, mitochondrial, and chloroplast proteins [23, 39, 40]. As a result, this PTM could be crucial for processes inside the nucleus and could be important for regulating different cellular and metabolic processes.

Proteins involved in photosynthesis are unique to plants and represent a notable proportion of the total observed Khib proteins. Of all the Khib proteins, 528 (49.2%) are located in the chloroplast (Supplemental Figure S3A), and 37 are involved in photosynthesis (Supplemental Figure S6; Table S8). This demonstrates that Khib may play a vital role in photosynthesis. For example, approximately 50% of the soluble leaf protein in C3 plants is ribulose-1,5-bisphosphate carboxylase (Rubisco), which is also the most abundant protein in plant leaves. In this study, the large subunit of Rubisco (RbcL, P11383) extensively consisted of Khib. This protein had up to 20 independent Khib residues, such as K175, K177, K201, K252, K227, K334, and K356 (Supplemental Table S1; Table 1). The IP and western blot assays using a pan-anti-Khib antibody verified that RbcL was indeed Khib (Figure 4A).

The majority of Khib sites occur in key function regions [46]. For example, the RbcL ε-amino groups, K175, K177, K201, and K252, play an important role in maintaining the stability of the -CO_2_-Mg^2+^ ternary complex [47]. This result implies that Khib is a key PTM in the regulation of photosynthetic CO_2_ assimilation and/or photorespiration in wheat. Moreover, the RbcL had overlapping key sites for Kac, Ksu, and Khib (Supplemental Figure S7). This indicates that Kac, Ksu, and Khib possibly compete or cooperate with each other in the regulation of the same protein. A study using *Arabidopsis* indicated that RbcL was a heavily acetylated protein [46, 47]. Previous study showed that de-acetylation of RbcL could increase the maximum enzyme catalytic activity by 40%, and this may further alter the photosynthetic rate [46]. Therefore, we speculate that the activity of RbcL could be simultaneously regulated by multiple PTMs. Moreover, three Khib sites (i.e., K32, K201, and K466) were found to be close to three phosphorylation sites (i.e., T34, S208, and T474) [48] (Supplemental Figure S7). As the phosphorylation is able to turn the corresponding protein functions on and off [39], we proposed that the interplay between Khib and nearby phosphorylation sites may result in the fine-tuning of protein functions. Other photosynthesis-related Khib proteins such as the small subunits of PS I and PS II, other subunit types of ATP synthase, and components of the photosynthetic electron transport chain were also found to be modified by Kac or Ksu in wheat [26, 27] (Supplemental Tables S12–S13). Thus, these PTMs may be highly dynamic and play a role in the regulation of photosynthesis. Future studies should examine the effects of Khib on photosynthesis, as well as aim to decipher how Khib coordinates phosphorylation, Kac, and Ksu in the regulation of Rubisco activity.

### Khib regulates protein functions

#### K206 is an important Khib site in PGK

In this study, many Khib metabolic enzymes were identified (Supplemental Table S1), which suggests that Khib may play a vital role in metabolic functions. For example, the majority of enzymes in tricarboxylic acid cycle (TCA), glycolysis/gluconeogenesis, and pentose phosphate pathways were modified by Khib. Altogether, 73 enzymes were determined to be Khib-modified in these three metabolic pathways; 16 of the 73 were abundantly modified and had over ten Khib sites (Supplemental Table S8). Ten enzymic reactions are required to convert glucose into pyruvate in glycolysis [49]; strikingly, seven of the key enzymes were highly modified by Khib, including GAPDH, pyruvate kinase (PKM), PGK, alpha-enolase (ENO), fructose-bisphosphate aldolase (FBA), triosephosphate isomerase (TPI), and fructose-1,6-bisphosphatase (FBP) (Supplemental Figure S8; Table S8). Among them, PGK catalyzes 3-phosphoglycerate (3-PG) and ATP to form 1,3-bisphosphoglycerate (1,3-BPG) and ADP, which is a reversible phosphotransfer reaction. The first ATP-yielding step of glycolysis is a PGK-catalyzed reaction that is crucial for energy generation [26]. The K206 site of PGK (W5H4V7) is critical for ATP binding and was modified by Khib. These results imply that the Khib PTM may be involved in the regulation of metabolic enzymes. Thus, Khib at this position likely affects protein function through some mechanism. However, future studies should examine the exact roles of Khib and its effects on proteins and other processes.

In this study, three PGK proteins (i.e., P12782, P12783/W5H4V7, and A0A1D6RDZ8) were determined to be Khib-modified. Among them, W5H4V7 was found to be modified by Khib at seven amino acid sites and was modified by Kma at multiple other sites [25–27] (**Figure 7A**). These proteins occurred in key functional regions (i.e., ATP binding and K206-hib) and shared the same lysine sites (K202-hib/ma) or were situated near the same functional regions (i.e., substrate binding, K35-hib, K58-hib, K75-ma, and K366-hib), implying that PGK may be functionally regulated by the Khib and Kma PTMs simultaneously. Additionally, the EMS-mutant wheat line with a premature stop codon in the PGK gene showed the delayed heading and stunted plants in comparison with wild-type controls (data not shown).

**Figure 7.**
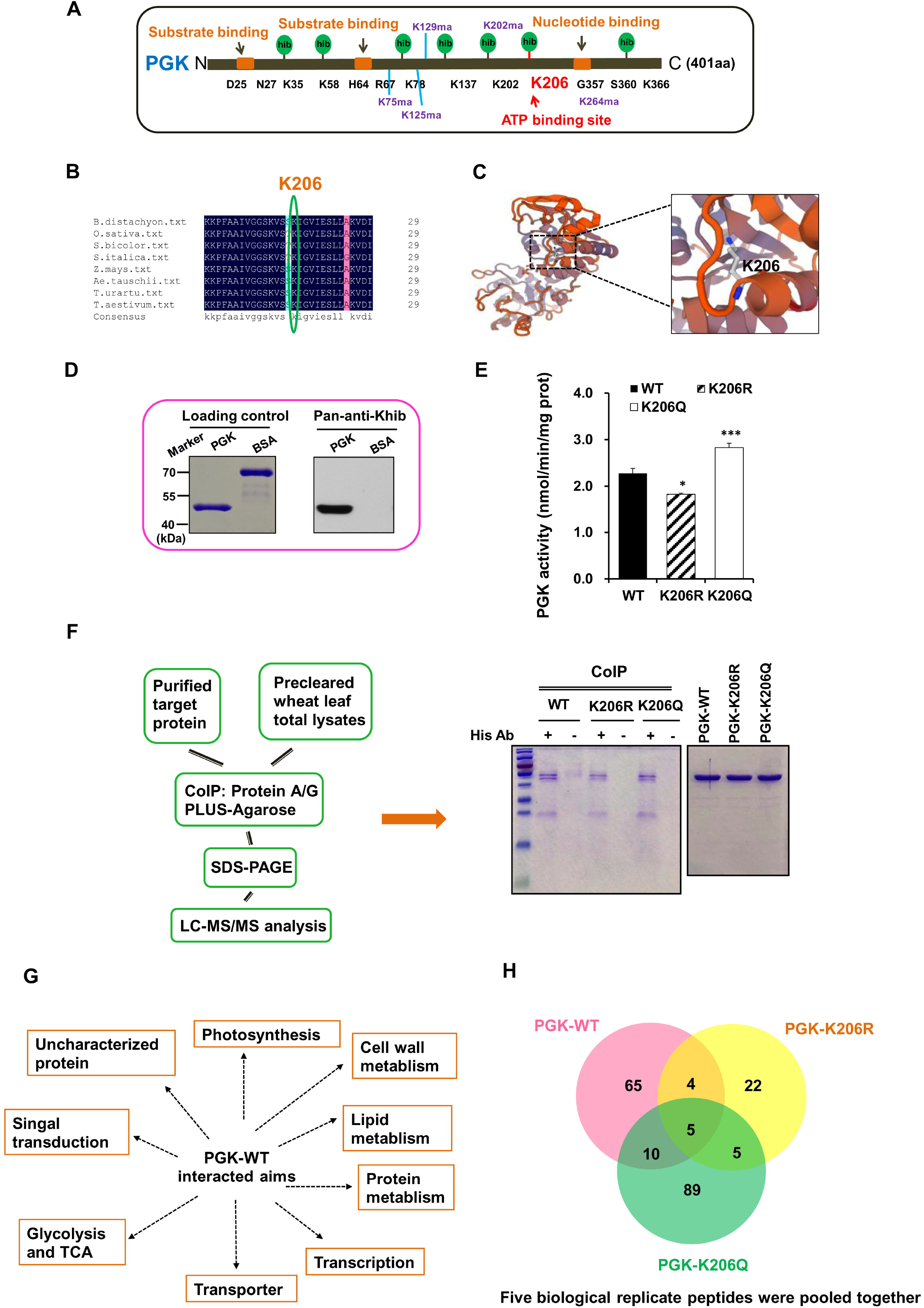
Khib regulates enzymatic functions of wheat PGK. (A) Location of lysine Kma and Khib sites. (B) Multiple sequence alignment of PKG from different species. (C) The tertiary structure prediction of PGK constructed by SWISS-MODEL. (D) Khib levels of PGK-WT and BSA were determined by SDS-PAGE (left) and western blot (right). (E) Effect of K206 Khib on enzymatic activity of PGK. Values are presented as the mean of three replicates. *: *P* < 0.05; ***: *P* < 0.001. (F) Schematic representation of PGK-WT and its mutants, and the strategy for identifying interacted proteins (left); Co-IP and LC-MS/MS analysis of PGK associated proteins (right). (G) Proteins interacted with PGK-WT. (H) Overlap of PGK-WT, PGK-206R, and PGK-206Q interacted proteins. Co-IP, co-immunoprecipitation; WT, wild type; Q, glutamine; R, arginine; LC-MS/MS, liquid chromatography and tandem mass spectroscopy.

Sequence alignment showed that the K206 site was evolutionarily conserved among all eight species investigated in this study (**Figure 7B**), and the K206 was modified only by Khib. The results of analyzing the protein secondary structures surrounding Khib sites revealed that four of the seven PGK Khib sites were located in *α*-helices, and three were located in disordered coils (Supplemental Figure S9). This indicates that the enzyme has no structural preference for either ordered secondary structures or disordered structures in Khib sites. Moreover, the K206 sites were located in α-helices that are known to be directly involved in ATP binding (**Figure 7C**). In addition to the mutagenesis experiment, the K206 site was mutated to both glutamine (K206Q, which is uncharged and could be used to simulate Khib) and arginine (K206R, which has a positive charge and cannot be modified by Khib) [35]. Western blot indicated that the PGK was indeed Khib (**Figure 7D**). We also assayed the enzymatic activity of the isolated proteins, which showed a significant decrease in enzymatic activity that resulted from the mutation of K206 to K206R. Interestingly, we found the mutation of K206 to K206Q could significantly increase the enzymatic activity as compared with the PGK-wild-type (WT) (**Figure 7E**). A previous study indicated that PGK activity was inhibited because of Kac on K220 disrupting the PGK binding with its ADP substrate in HEK293T cells [50]. Thus, we speculate that Khib at K206 could directly perturb the ATP binding site.

To further examine whether the ATP binding site mutation affects the interaction function of PGK, precleared wheat leaf proteins and PGK-WT, PGK-K206R, or PGK-K206Q were immunoprecipitated using beads **(Figure 7F**). Interactive proteins were co-precipitated with the PGK-WT protein or its two mutants and were then eluted and visualized by Coomassie blue staining. Experiments were repeated independently five times. Results revealed that the PGK-WT interacted with 84 proteins involved in multiple metabolic pathways (**Figure 7G**; Supplemental Table S14). However, 35 proteins were found to interact with the PGK-K206R, and 109 proteins interacted with the PGK-K206Q (**Figure 7H**; Supplemental Table S14), which is consistent with enzymatic activity levels. Among these proteins, five were common in the PGK-WT and its two mutants (**Figure 7H**). These results suggest that K206 was important for PGK enzymatic activity and that Khib was likely affecting enzymatic activity. However, these lysine sites may also be involved with the various other PTMs existing in the organism.

#### Khib regulates 14-3-3 protein interaction

Numerous biological processes and pathways depend on the 14-3-3 interactions to regulate key metabolic points [51]. The 14-3-3s proteins could bind to specific phosphorylated motifs of target proteins by forming homodimers and heterodimers in the native state [52]. To understand the role of 14-3-3 Khib and the impact of Khib on the interaction of 14-3-3 with proteins or phosphorylated proteins in wheat, three Khib lysines (K56, K124, and K129) that were highly conserved among 14-3-3 proteins were mutated (Supplemental Figure 10A–B). Khib lysines of 14-3-3 proteins were mutated into the R and Q forms. Interactive proteins were co-precipitated with a 14-3-3 protein or its 6 mutants before being eluted and visualized by Coomassie blue staining (Supplemental Figure 10C). The analysis showed the 14-3-3 WT interacts with 241 proteins involved in multiple metabolic pathways (Supplemental Table S15). In total, 321 and 302 proteins were found to interact with 14-3-3 K56R and K56Q, respectively (Supplemental Table S15). More than two-fold (586, 585, 551, and 546) proteins were identified to interact with double mutants (14-3-3 K124+129R and K124+129Q) or triple mutants (K56+124+129R and K56+124+129Q) when compared with WT (Supplemental Table S15). Compared with WT (with 27, the more phosphorylated proteins (34, 32, 94, 98, 85, and 79) were found to interact with the mutants, especially with double and triple mutants (Supplemental Table S16). The phosphopeptides identified by LC-MS/MS contained sequence-specific motifs, including RSxpSxP, RSxxpSxP, and YpT [52]. Our data indicated that phosphorylation-dependent interactions existed on the phosphopeptide-binding domain, implying that both of phosphorylation and Khib may have influence on the function of 14-3-3.

### Wheat sirtuin and Zn-binding HDAC proteins regulate Khib state

Debutyrylase and decrotonylase activity has been observed in mammalian Zn-finger HDAC proteins [39]. For these activities, TSA is an inhibitor of HDAC I and II [53], and NAM is an inhibitor for SIRT family deacetylases [54, 55]. NAM- and TSA-treated wheat seedlings were investigated to determine if wheat leaf Khib could be regulated by SIRT or Zn-binding HDAC proteins; this was observed by immunoblots using the pan-anti-Khib antibody and pan-anti-Kac antibody. Compared with Kac (**Figure 8A**), we found that the Khib level in wheat seedlings did not increase by much after TSA and NAM treatments in the proteins with molecular weights greater than 25 kDa; however, the Khib level increased in the low-molecular-weight proteins (<25 kDa) (**Figure 8B**), suggesting that wheat SIRT or Zn-binding HDAC proteins regulated Khib. To date, the preferences and targets of the different HDACs in plants are still unknown. Compared with the genomes of *Arabidopsis* (125 Mb) [56] and rice (466 Mb) [57], common wheat possesses a larger genome (16 Gb) and is an allohexaploid (AABBDD) [58]. It has been proven that at least 18 HDACs from the three different families (RPD3/HDA1, SIRT2, and HD families) were encoded in *Arabidopsis* [59] and rice [60]. Hexaploidy wheat possibly contains more HDACs, meaning that studies on the impact of HDAC-regulation of Khib proteins in *Triticum aestivum* L are more difficult than in other, smaller-genome species.

**Figure 8.**
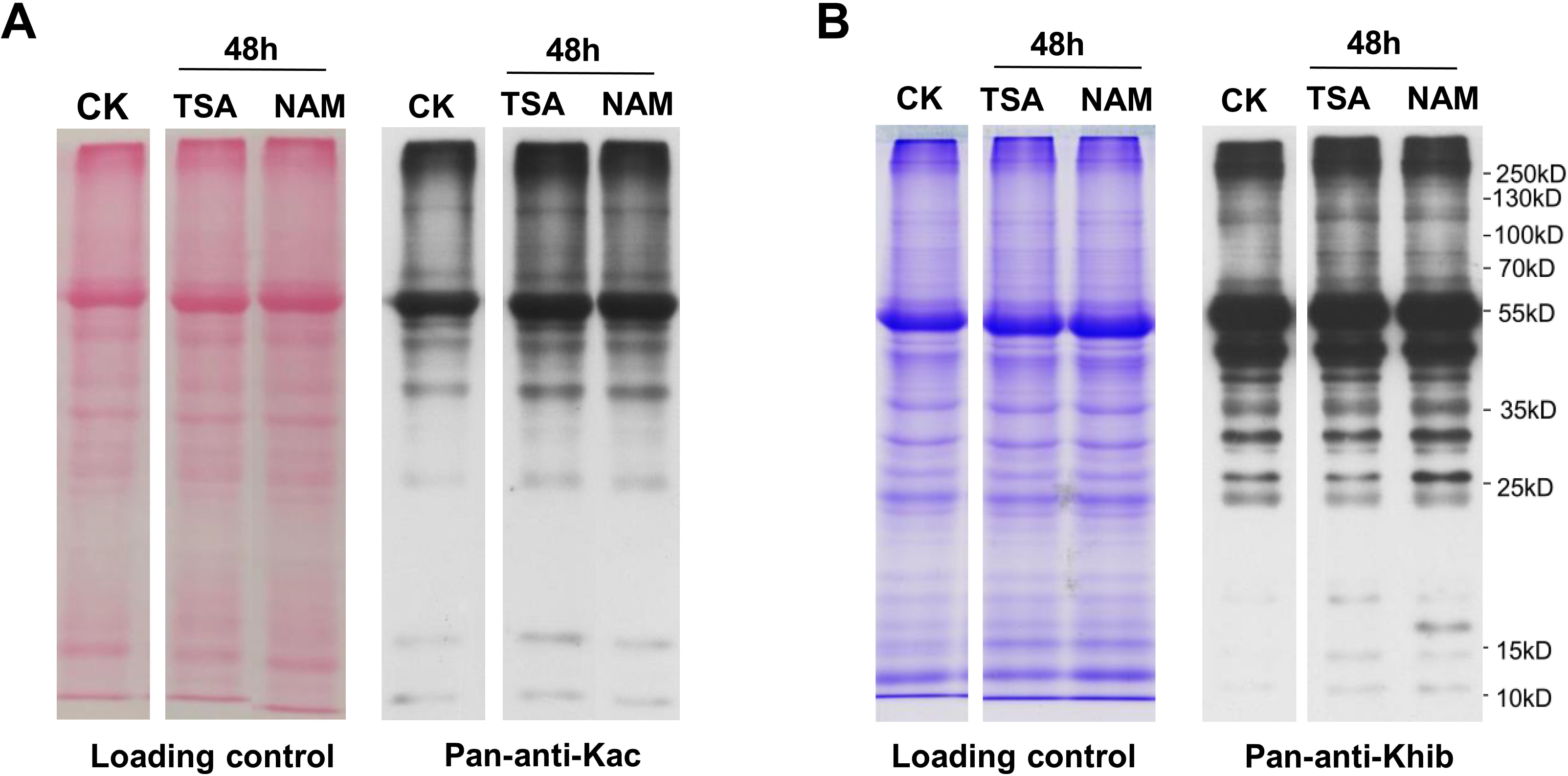
TSA and NAM treatments change wheat Khib state. CK, control; TSA, trichostatin; NAM, nicotinamide; pan-anti-Kac antibody, pan-anti-acetyllysine antibody.

## Conclusions

In this study, we confirmed that Khib is an evolutionarily-conserved PTM in wheat and its donor progenitors. Based on affinity purification and high-resolution LC-MS/MS, a global 2-hydroxyisobutyrylome was performed in common wheat for the first time, and a total of 3348 Khib sites in 1047 proteins were identified. Analysis showed that Khib was found to participate in many kinds of biological processes, such as photosynthesis, glycolysis, and protein metabolism. IP and western blot confirmed that Khib proteins had an *in vivo* origin. Mutagenesis experiments demonstrated that Khib on the K206 site was a key regulatory modification of PGK activity and affected protein interactions. Moreover, western blot indicated that wheat sirtuin or Zn-binding HDAC proteins mainly regulated the low-molecular-weight Khib proteins. This work improves our understanding of this novel lysine PTM, and the large set of Khib proteins identified in common wheat will act as an important data source for future functional studies in plants.

## Materials and methods

### Plant materials and growth conditions

Wheat donor progenitors, including *Triticum urartu* (PI428185) with an AA genome, *Aegilops speltoides* (PI542245) with an SS genome, *Aegilops tauschii var. strangulate* (AS2393) with a DD genome, tetraploid *Triticum durum* Langdon (AABB), and hexaploid wheat Aikang 58 (Ak58), were planted as detailed in our previous report [42]. Leaf tissues from two seedling leaf periods were collected. Afterward, samples were frozen in liquid nitrogen for subsequent protein extraction and experimentation.

### Protocol for 2-hydroxyisobutyrylome

Leaf tissues collected from two seedling leaf periods were used to extract proteins (Supplemental Method S1). Protein concentrations were examined according to the manufacturer’s instructions of the BCA Protein Assay Kit (Beyotime, China, Catalog No. P0012). Trypsin digestion, HPLC fractionation, affinity enrichment, LC-MS/MS analysis, and database searches are shown in Supplemental Method S1. We have deposited mass spectrometry proteomics data to the ProteomeXchange Consortium via the PRIDE [61] partner repository with the dataset identifier PXD012666.

### Bioinformatics methods

UniProt-GOA database (http://www.ebi.ac.uk/GOA/) was used to carry out the GO annotation. InterProScan was used to annotate proteins’ domain functional descriptions. Protein pathways were annotated using the KEGG database. Wolfpsort was used to predict subcellular localization, and Motif-X was used to analyze the model of sequences. Genes/Proteins (STRING) database v10.5 was used for determining protein-protein interactions; the interaction network derived from STRING was visualized in Cytoscape [62].

### IP assay

Wheat leaf (0.5 g) proteins were lysed in lysis buffer [1% Triton X-100, 20 mM Tris-HCl (pH 8.0), 2 mM DTT, 800 μM PMSF, 250 mM sucrose], consisting of a protease inhibitor cocktail. IP was carried out by incubating with or without 25 μL of anti-RbcL (Real-Time Biotechnology, Catalog No. RGR2030). Thirty microliter of Protein A/G PLUS-Agarose (Santa Cruz Biotechnology, USA, Catalog No. sc-2003) was added to the solution and incubated for 6 h on ice. After incubation, we washed the bound proteins four times with phosphate buffer solusion (PBS) buffer, then suspended them in 40 μL of PBS buffer. Proteins in PBS buffer were analyzed by sodium dodecyl sulfate-polyacrylamide gel electrophoreses (SDS-PAGE) and western blot.

### Wheat germ protein expression system

Transcripts of CAT (A0A1D5YMQ8), 14-3-3 (L0GED8), GST (Q9SP56), GAPDH (A0A1D5TTT4), and PGK (W5H4V7) were amplified by PCR from the cDNA of wheat Ak58 (Supplemental Table S17). After restriction digestion, CAT, 14-3-3, GST, PGK, and GAPDH were cloned into pFN19K HaloTag® T7 SP6 Flexi® Vector (Promega, USA, Catalog No G184A) with Sgf I and Pme I restriction enzyme cutting sites; a 6xHis tag was added before TAG. The recombinant Halo-His-gene was expressed using a 30 μL TNT®SP6 High-Yield Wheat Germ Protein Expression System (Promega, USA, Catalog No L3261). Expression levels were determined by SDS-PAGE and western blot. Magne® Halo Tag® Beads (Promega, USA, Catalog No G728A) were used to purify recombinant proteins according to the manufacturer’s instructions. We used Halo TEV protease (Promega, USA, Catalog No. L3261) to cleave the Halo tag and then captured the recombinant His-protein in elution buffer (137 mM NaCl, 2mM KH_2_PO_4_, 10 mM Na_2_HPO_4_, 2.7 mM KCl, and 0.005% IGEPALR CA-630). Lastly, we analyzed the recombinant proteins by SDS-PAGE and western blot.

### Site-directed mutagenesis and prokaryotic expression

PGK (W5H4V7) was cloned into the prokaryotic expression vector, PET28a, using NdeI and XhoI as restriction enzyme cutting sites (Supplemental Table S17). We performed site-directed mutagenesis of K206 (K206R, AAG-AGG; K206Q, AAA-CAA) within the PET28a plasmid using the Fast Site-Directed Mutagenesis Kit (Tiangen, China, Catalog No. KM101). PCR was conducted using site-specific primers (Supplemental Table S17). Positive mutants were verified by DNA sequencing. *E. coli* BL21 (DE3) was used to express recombinant PGK WT, PGK-K206R, and PGK-K206Q. A His-tag Protein Purification Kit (Beyotime, China, Catalog No. P2226) was used to purify proteins. The collected PGK recombinant proteins and two mutants were analyzed by SDS-PAGE and western blot. For PGK, enzyme activity was measured by following the decrease in absorbance at 340 nm, which is due to the oxidation of NADH. The assay was conducted using commercial assay kits (Comin Biotechnology, Suzhou, China) according to the manufacturer’s instructions. Experiments were performed in triplicate.

14-3-3 was also cloned into the prokaryotic expression vector PGEX-6P-1 using BamHI and NotI as restriction enzyme cutting sites (Supplemental Table S17). Site-directed mutagenesis of K56 (K56R, AAA-AGA; K56Q, AAA-CAA), K124+129 (K124+129R, AAA-AGA; AAA-AGA; K124+129Q, AAA-CAA, AAA-CAA), and K56+124+129 (K56+124+129R, AAA-AGA, AAA-AGA, AAA-AGA; K56+124+129Q, AAA-CAA, AAA-CAA, AAA-CAA) within the PGEX-6P-1 plasmid was performed using the fast Site-Directed Mutagenesis Kit (Tiangen, China, Catalog No. KM101). PCR was conducted using site-specific primers (Supplemental Table S17). Positive mutants were verified by DNA sequencing. *E*. *coli* BL21 (DE3) was used to express recombinant 14-3-3-WT, K56R, K56Q, K124+129R, K124+129Q, K56+124+129R, and K56+124+129R proteins. A glutathione S-transferase (GST)-tag Protein Purification Kit (Beyotime, China, Catalog No P2262) was used to purify proteins. We analyzed the collected 14-3-3 recombinant proteins and six mutants by SDS-PAGE and western blot.

### Co-immunoprecipitation of wheat leaf proteins with PGK/14-3-3 and its mutants

Wheat leaf extracts were precleared per the following method. Approximately 1 mg of whole wheat leaf extract (in line with IP) was added to 30 μL of suspended agarose conjugate (Protein A/G PLUS-Agarose: sc-2003) brought to 1 mL in volume with PBS buffer. This solution was incubated at 4 °C for 30 min, then centrifuged with pellet beads at 3,000 rpm for 30 s at 4 °C. The supernatant was transferred to a new microcentrifuge tube and kept at 4 °C. Then, 200 μg of PGK, PGK-206R, or PGK-206Q proteins and 25 μL of His-tag Mouse monoclonal antibody (Abmart, China, Catalog No M30111) were added and incubated at 4 °C overnight with mixing. After incubation, the mix was washed four times with PBS buffer, then suspended in 40 μL PBS buffer, and separated by SDS-PAGE gel. Based on molecular weight, we cut each lane into 2 or 3 segments. Trypsin was used to digest gel segments. Five biological replicate peptides of PGK-WT, PGK-206R, or PGK-206Q were pooled together. Pooled tryptic peptides were used for LC-MS/MS analysis. Co-immunoprecipitation (Co-IP) of the wheat leaf proteins with 14-3-3 and its six mutants was conducted using the method detailed above.

### Nicotinamide- and trichostatin A-treated seedlings

Wheat Ak58 seedlings from the two leaf periods were treated with 0.5 μM trichostatin A (TSA) (MedChemExpress, USA, Catalog No. HY-15144) or 10 mM nicotinamide (NAM) (MedChemExpress, USA, Catalog No. HY-B0150) for 48 h. Leaf tissues from WT, TSA-treated, and NAM-treated seedlings were collected and then frozen in liquid nitrogen for subsequent protein extraction and experimentation.

### Antibodies

The antibodies, anti-RbcL (Real-Time Biotechnology, China, Catalog No. RGR2030), plant Actin monoclonal (Abbkine, USA, Catalog No. A01050), anti-HaloTag monoclonal (Promega, USA, Catalog No. G921A), anti-His-tag mouse monoclonal (Abmart, China, Catalog No. M30111), anti-GST-tag mouse monoclonal (Abmart, China, Catalog No. M20007), pan-anti-Khib (PTM Biolabs, China, Catalog No. PTM-801), and pan-anti-acetyllysine (pan-anti-Kac, PTM Biolabs, China, Catalog No. PTM-101) were used in this study. We also used goat anti-mouse lgG HRP (Abmart, China, Catalog No. M21001), goat anti-rabbit lgG HRP (Abmart, China, Catalog No. M21002), and HRP-conjugated AffiniPure mouse anti-rabbit IgG light chain (ABclonal, USA, Catalog No. AS061) as secondary antibodies in this study.

### Statistical analyses

A one-way analysis of variance (ANOVA) and Duncan’s multiple range test (DMRT) were carried out using SPSS v17.0 statistical software.

## Data availability

The mass spectrometry proteomics data in this study have been deposited in the ProteomeXchange Consortium via the PRIDE [61] partner repository with the dataset identifier PXD012666. http://www.ebi.ac.uk/pride.

## Author contributions

FC and NZ designed and supervised the project. NZ, LZ (Lingran Zhang), and LL performed vector construction, protein purification, and western blot assays. NZ and LL performed IP and CoIP assays. LZ (Lingran Zhang), JG, and LZ (Lei Zhao) performed site-directed mutagenesis assays. NZ participated in the sequence alignment. NZ, YR, ZD, and FC performed the proteomic data analyses. NZ and FC wrote the paper. All authors have read and approved the final manuscript.

## Competing interests

The authors have declared no competing interests.

## Acknowledgments

This project was funded by the National Key Research and Development Program (2016YFD0101802), the National Natural Science Foundation (U1804234, 31901542, and 3181101544), the Henan Natural Science Foundation (182300410022), the Henan Science and Technology Foundation (182102110353), and the Science and Technology Innovation Foundation of Henan Agricultural University (KJCX2018A01) in China. We thank Prof. Jirui Wang from Sichuan Agricultural University for kindly providing wheat donor progenitors, and Jingjie PTM BioLab (Hangzhou, China) and Shanghai Applied Protein Technology for technical support.

## Supplemental materials

**Table S1.** Quantitative results of lysine 2-hydroxyisobutyrylation sites identified in common wheat.

**Table S2.** Structural features analysis of 2-hydroxyisobutyrylation proteins in common wheat.

**Table S3.** Subcellular location of 2-hydroxyisobutyrylation proteins in common wheat.

**Table S4.** GO annotation 2-hydroxyisobutyrylation proteins in common wheat.

**Table S5.** GO enrichment of 2-hydroxyisobutyrylation proteins in common wheat.

**Table S6.** KEGG enrichment of 2-hydroxyisobutyrylation proteins in common wheat.

**Table S7.** Protein domain functional description of 2-hydroxyisobutyrylation proteins in common wheat.

**Table S8.** KEGG pathways of 2-hydroxyisobutyrylation proteins in common wheat.

**Table S9.** Information of the interaction network among 2-hydroxyisobutyrylation proteins in common wheat using STRING.

**Table S10.** Overlap of the Khib proteins among common wheat, rice, and *Physcomitrella patens*.

**Table S11.** KEGG enrichment of common 2-hydroxyisobutyrylation proteins identified in wheat, rice, and *Physcomitrella patens*, and specific 2-hydroxyisobutyrylation proteins identified in wheat.

**Table S12.** Overlap of wheat 2-hydroxyisobutyrylation, malonylation, succinylation, and acetylation proteins.

**Table S13.** Overlap of wheat 2-hydroxyisobutyrylation, malonylation, succinylation, and acetylation sites.

**Table S14.** LC-MS/MS analysis of PGK-WT, PGK-K206R, and PGK-K206Q associated proteins.

**Table S15.** LC-MS/MS analysis of 14-3-3-WT, 14-3-3-K56R, 14-3-3-K56Q, 14-3-3-K124+129R, 14-3-3-K124+129Q, 14-3-3-K56+124+129R, and 14-3-3-K56+124+129Q associated proteins.

**Table S16.** Phosphorylation LC-MS/MS analysis of 14-3-3-WT, 14-3-3-K56R, 14-3-3-K56Q, 14-3-3-K124+129R, 14-3-3-K124+129Q, 14-3-3-K56+124+129R, and 14-3-3-K56+124+129Q associated proteins.

**Table S17.** Primer sequences used in this study.

**Method S1.** The procedure of plant protein extract, trypsin digestion, HPLC fractionation, affinity enrichment, LC-MS/MS analysis, and database search.

**Figure S1.** Examples of raw mass spectra in common wheat.

**Figure S2.** Mass error distribution of identified peptide in common wheat.

**Figure S3.** Pie charts of the distribution of 2-hydroxyisobutyrylation proteins based on their predicted subcellular localization (A), cellular components (B), molecular functions (C), and biological processes (D).

**Figure S4.** Interaction networks of the 2-hydroxyisobutyrylation proteins in wheat using String software.

**Figure S5.** The tertiary structure prediction information of 14-3-3 (L0GED8) using SWISS-MODEL.

**Figure S6.** 2-hydroxyisobutyrylation proteins involved in photosynthesis.

**Figure S7.** Location of lysine, acetylation, succinylation, malonylation, 2-hydroxyisobutyrylation, and phosphorylation sites in large subunit of Rubisco (RbcL, P11383) of wheat.

**Figure S8.** 2-hydroxyisobutyrylation proteins involved in glycolysis/gluconeogenesis.

**Figure S9.** The secondary structures analysis of phosphoglycerate kinase (PGK, W5H4V7).

**Figure S10.** Effects of Khib on 14-3-3 with its interactive proteins.

