## Supplemental Figure S1-S10 for "Global Profiling of 2-hydroxyisobutyrylome in Common Wheat"

**Supplemental Figure S1. Examples of raw mass spectra in common wheat.**

**A0A0C4BJE5**

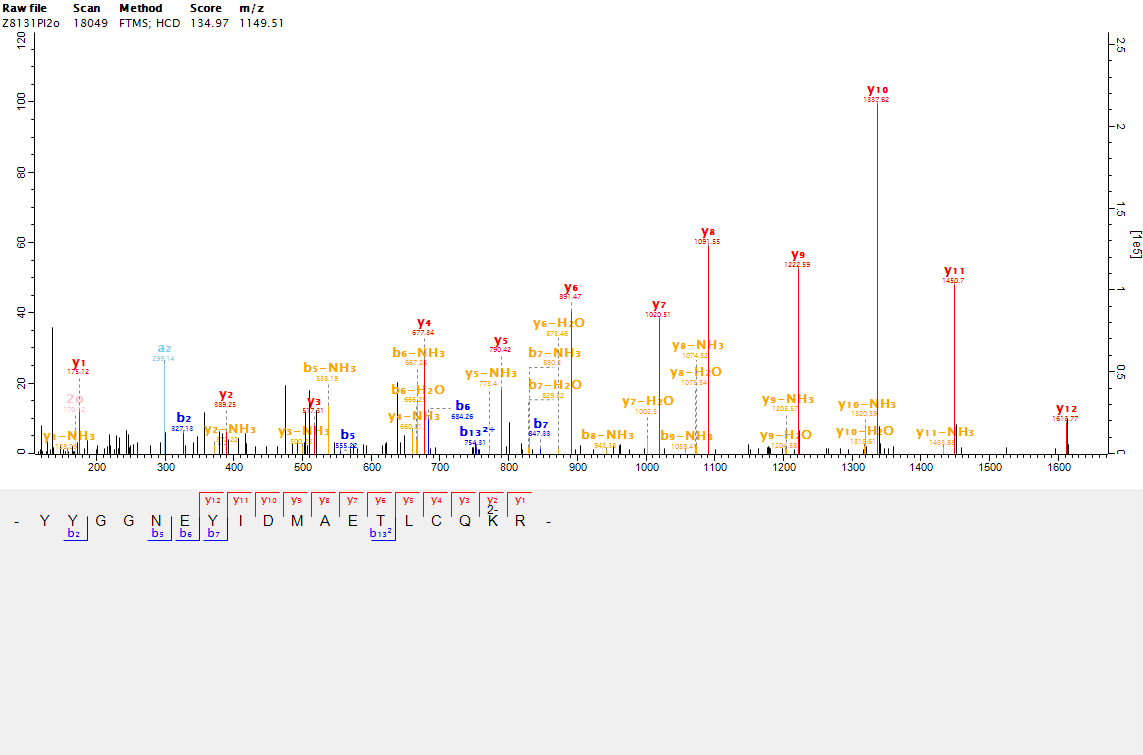

**A0A1D5RZX1**

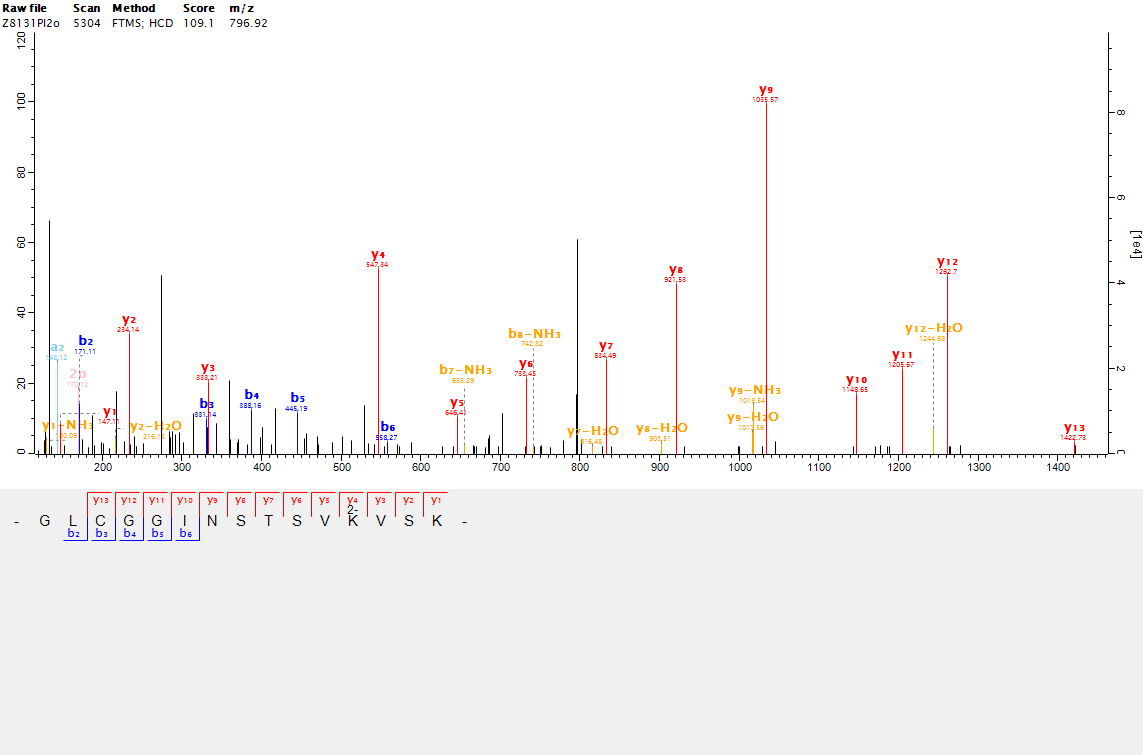

**A0A1D5SSJ7**

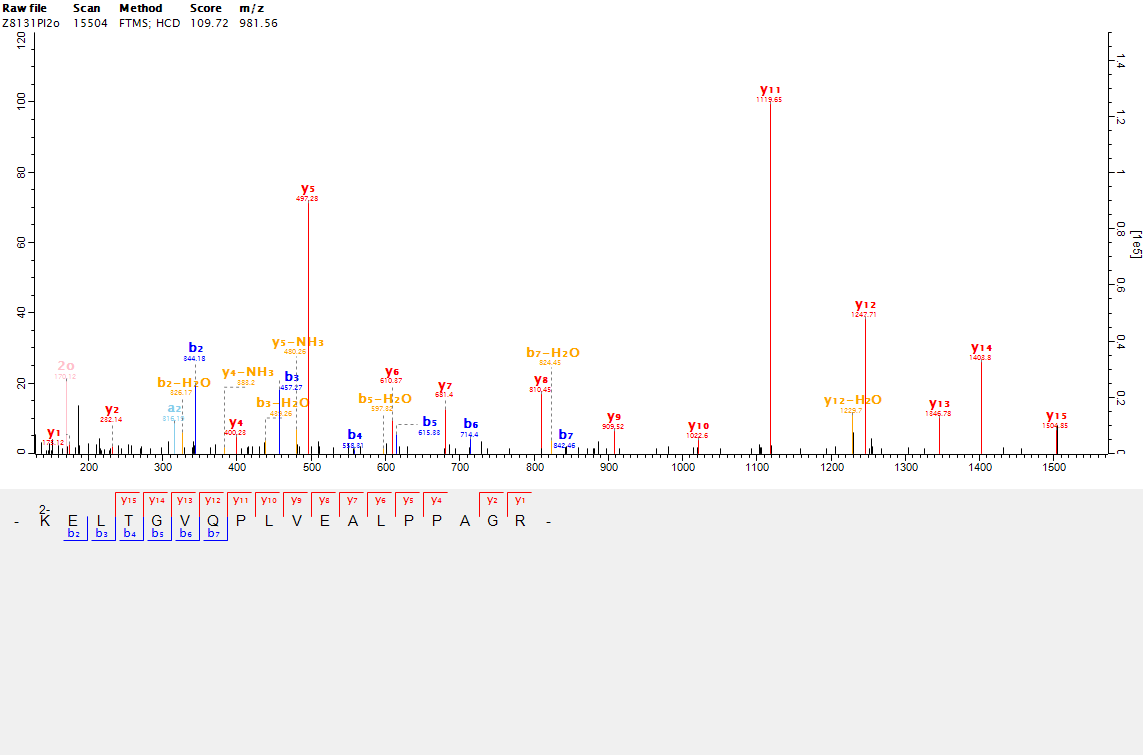

**F4Y593**

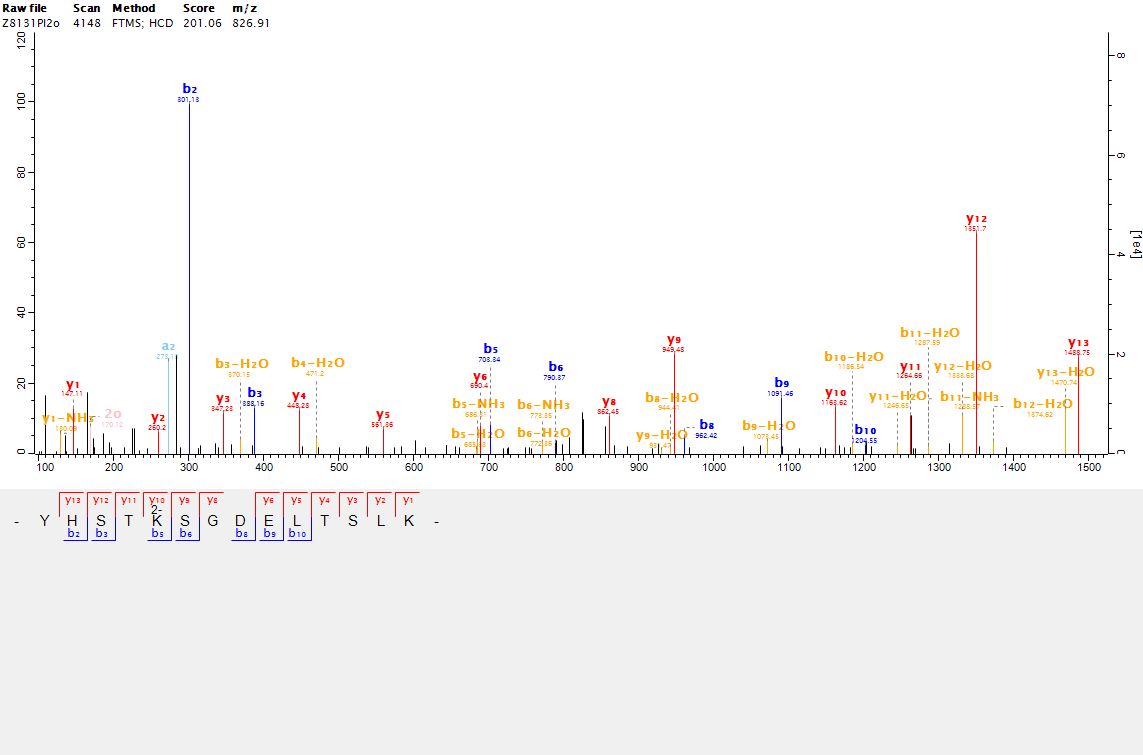

**P05151**

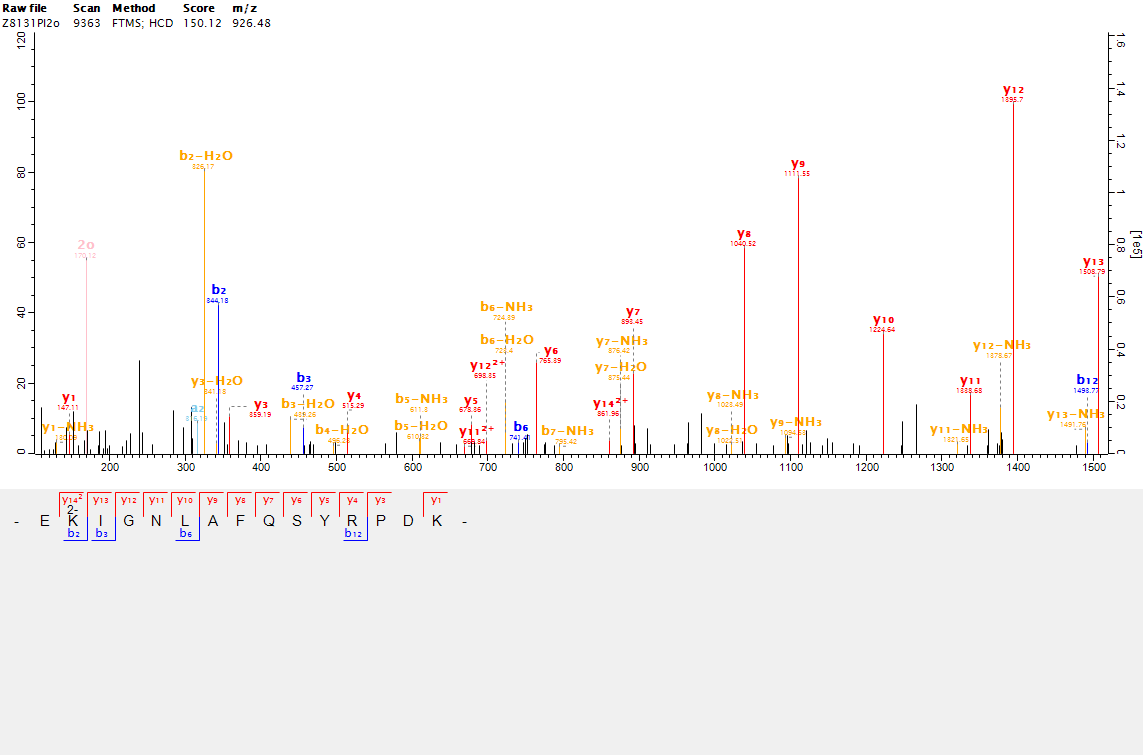

**Q9SP56**

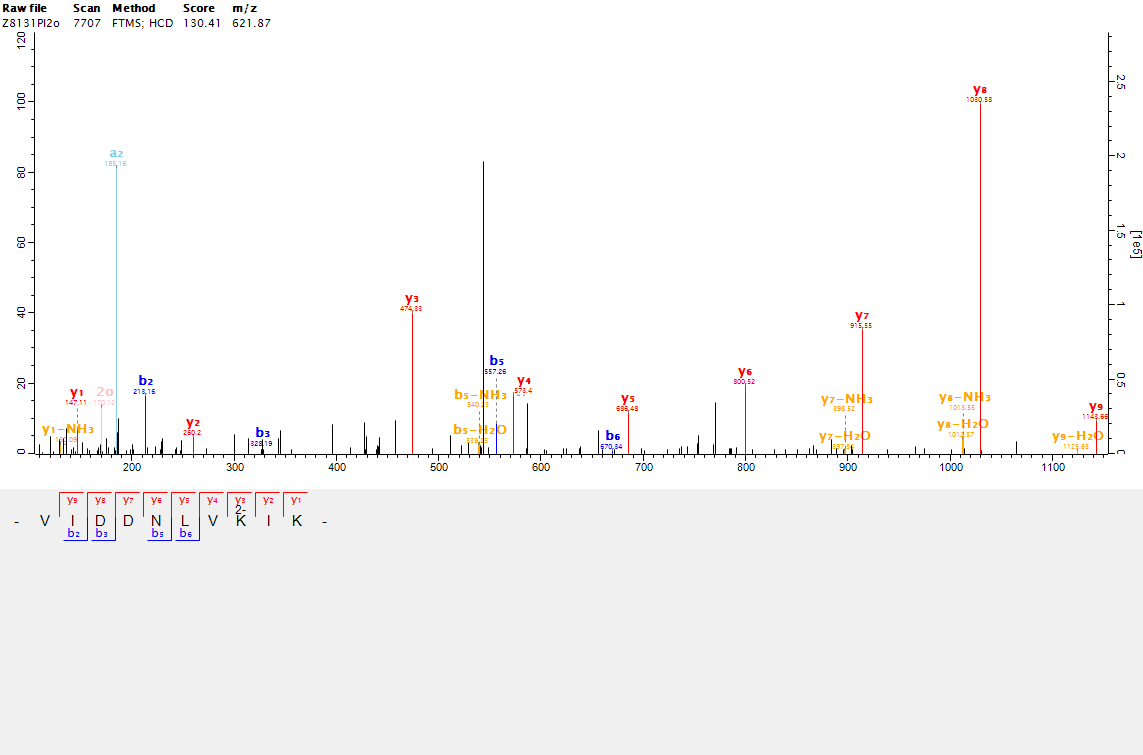

**W4ZSC8**

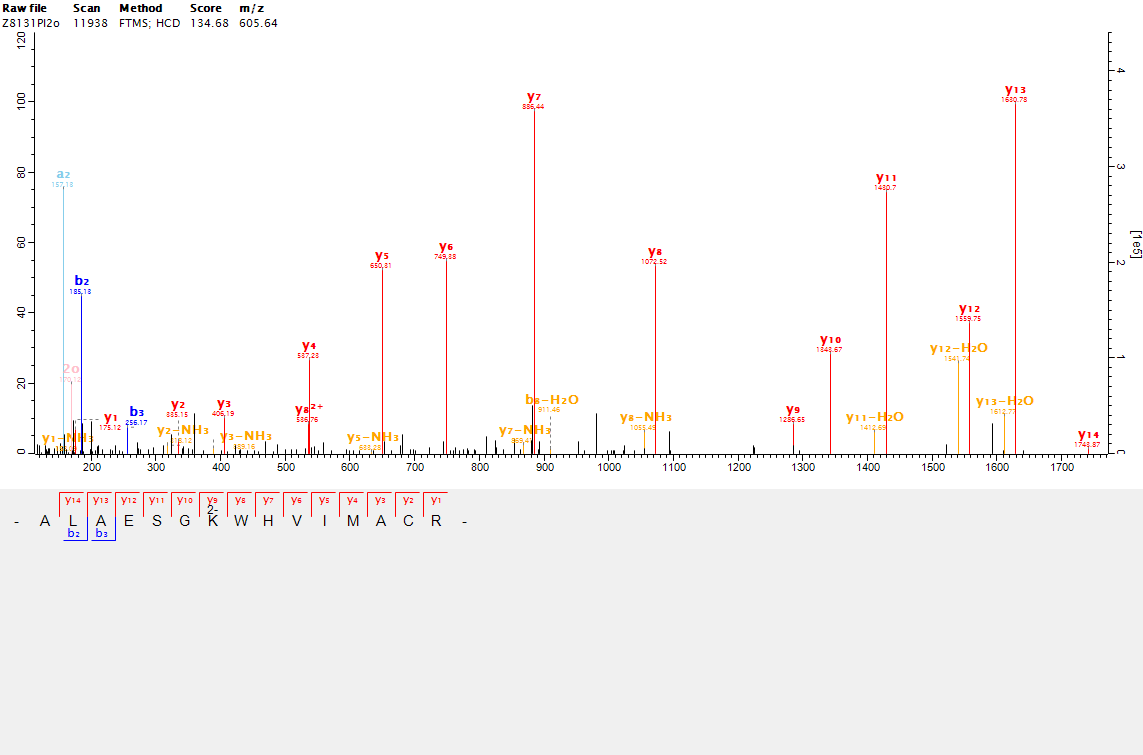

**W5C4P1**

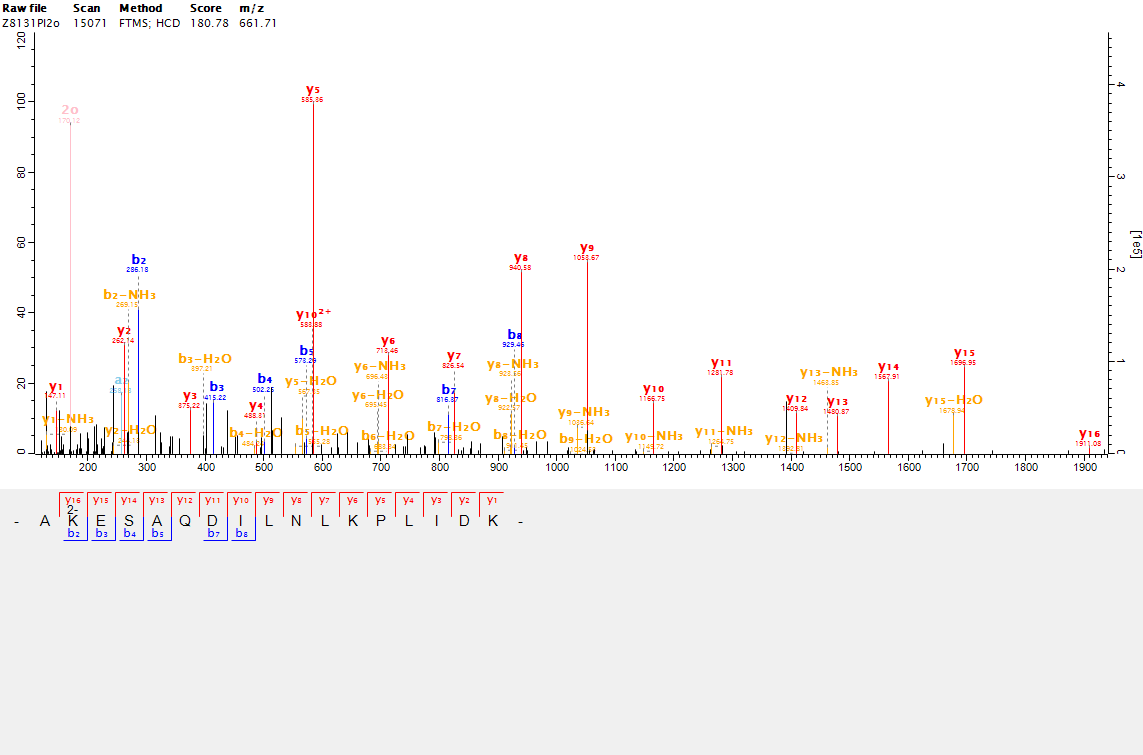

**W5FEE5**

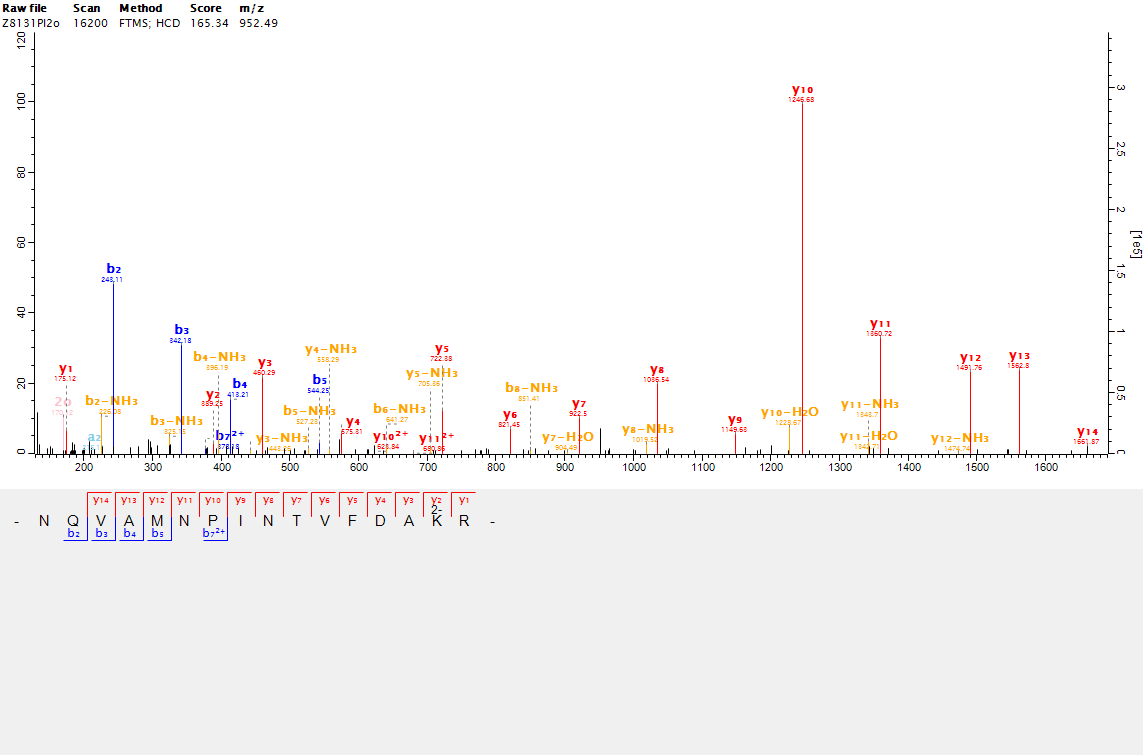

**W5GIE6**

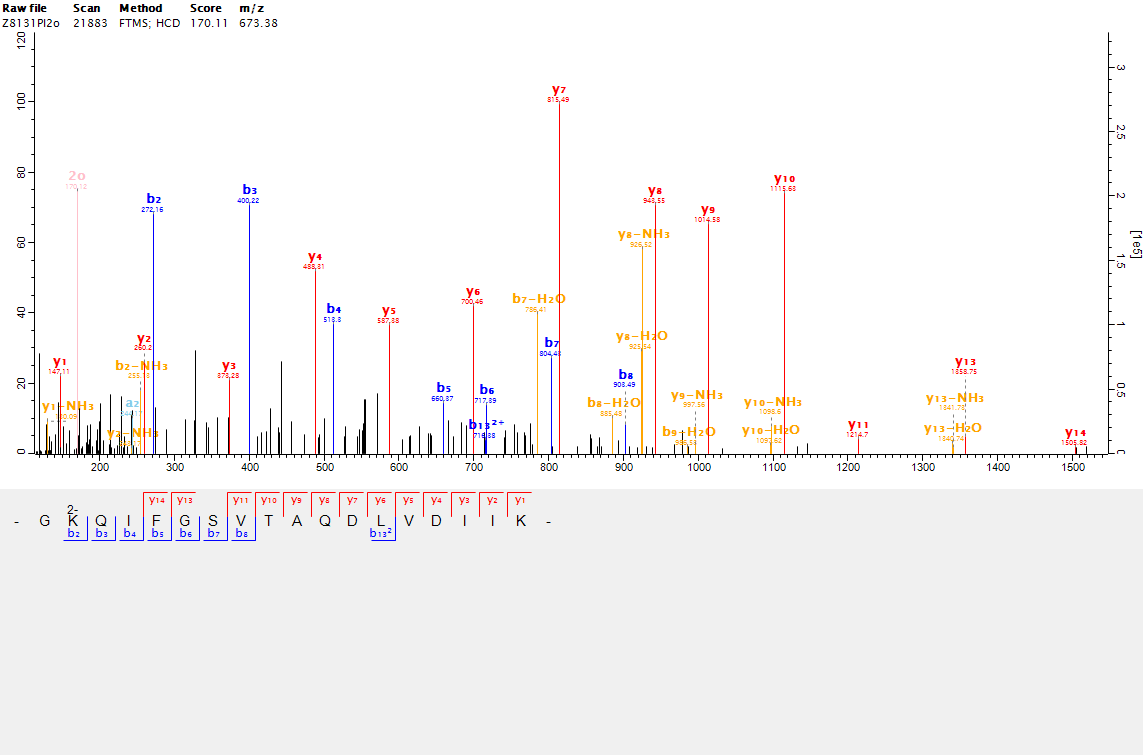

**Supplemental Figure S2. Mass error distribution of identified peptide in common wheat.**

| Peptide mass error [ppm] | Peptide score |
| --- | --- |
|  | 65.03 |
| -2.185 | 196.67 |
| -2.5218 | 105.52 |
| -2.2395 | 152.07 |
| -1.5485 | 134.61 |
| -1.6279 | 114.64 |
| -1.7267 | 126.83 |
| -1.6104 | 174.19 |
| -1.7061 | 111.66 |
| -1.5374 | 83.758 |
| -2.021 | 117.2 |
| -1.585 | 87.802 |
| -2.0302 | 118.72 |
| -0.97282 | 98.654 |
| -2.24 | 53.033 |
| -1.938 | 125.03 |
| -1.7779 | 131.82 |
| -1.5717 | 140.78 |
| -1.9858 | 150.35 |
|  | 82.069 |
| -1.8333 | 89.827 |
| -1.5759 | 50.271 |
| -1.6655 | 109.39 |
| -1.5184 | 70.225 |
| -1.4347 | 95.273 |
|  | 96.229 |
| -1.8163 | 111.61 |
| -2.9102 | 72.789 |
| -2.2763 | 101.17 |
| -1.9963 | 168.64 |
| -1.6901 | 80.412 |
| -2.1844 | 121.68 |
| -1.3809 | 108.71 |
| -1.1859 | 167.97 |
| -1.6974 | 97.813 |
| -2.0098 | 88.108 |
| -1.3682 | 193.77 |
| -1.8985 | 167.29 |
|  | 72.891 |
|  | 50.077 |
| -2.1457 | 80.316 |
| -2.1667 | 129.83 |
| -1.7291 | 139.48 |
| -0.99178 | 100.7 |
| -1.3768 | 152.62 |
| -2.5243 | 65.085 |
| -2.2358 | 247.29 |
| -2.5668 | 63.419 |
| -1.4856 | 77.24 |
| -1.4571 | 98.811 |
| -1.5402 | 153.18 |
| -2.0333 | 150.91 |
| -1.6406 | 115.71 |
| -1.7713 | 107.03 |
| -1.681 | 214.26 |
|  | 61.127 |
| -2.2525 | 223.31 |
| -2.2485 | 49.333 |
| -2.0637 | 98.407 |
| -1.762 | 66.809 |
| -1.4985 | 90.385 |
|  | 91.626 |
| -0.89083 | 60.541 |
| -2.1428 | 138.85 |
| -1.5292 | 69.331 |
| -1.7898 | 57.434 |
|  | 53.404 |
| -2.1442 | 126.71 |
| -1.7207 | 152.88 |
| -1.75 | 175.38 |
| -2.0623 | 48.044 |
| -1.9269 | 70.837 |
| -2.1428 | 138.66 |
| -1.8224 | 78.078 |
| -1.9623 | 120.65 |
| -2.2976 | 139.34 |
| -1.8057 | 167.32 |
| -2.031 | 106.6 |
| -1.8093 | 78.324 |
| -1.7601 | 85.287 |
|  | 44.054 |
| -2.1779 | 121.9 |
| -2.194 | 84.878 |
| -2.0512 | 128.06 |
| -1.3208 | 74.657 |
| -2.1751 | 111.95 |
| -2.2276 | 139.7 |
|  | 62.68 |
| -1.7553 | 88.243 |
| -2.1805 | 155.66 |
| -2.0457 | 123.02 |
| -2.0238 | 147.12 |
|  | 111.82 |
| -2.0459 | 105.52 |
| -1.7243 | 71.879 |
| -1.7048 | 81.877 |
| -1.7945 | 157.86 |
|  | 92.445 |
|  | 93.096 |
| -1.6299 | 125.97 |
| -1.7595 | 145.69 |
| -1.3061 | 117.2 |
| -1.3254 | 78.326 |
| -2.0284 | 96.067 |
| -2.0393 | 107.03 |
|  | 61.127 |
| -1.6718 | 57.267 |
| -2.3906 | 84.817 |
| -2.2072 | 77.662 |
| -1.3391 | 71.601 |
| -2.215 | 76.586 |
| -2.044 | 91.095 |
| -2.6123 | 63.419 |
| -1.7893 | 98.898 |
| -2.8518 | 121.68 |
| -1.7171 | 78.191 |
| -2.2218 | 86.539 |
| -1.3362 | 88.427 |
| -2.0425 | 136.33 |
| -1.9645 | 197.75 |
| -1.3042 | 110.31 |
| -1.8813 | 111.82 |
| -2.0813 | 200.09 |
| -1.5911 | 98.804 |
| -1.7063 | 155.88 |
|  | 49.298 |
| -2.4227 | 120.87 |
| -2.2648 | 92.865 |
| -1.888 | 99.834 |
| -1.6306 | 48.354 |
| -1.7302 | 84.753 |
| -1.2002 | 73.273 |
|  | 55.437 |
| -2.3552 | 75.189 |
| -1.6595 | 46.171 |
| -1.0728 | 172.43 |
| -1.3294 | 56.174 |
| -2.1754 | 58.246 |
| -1.8596 | 147.1 |
| -1.8401 | 125.75 |
| -1.9667 | 131.72 |
| -2.1498 | 95.531 |
| -2.3253 | 103.55 |
| -2.3965 | 113.89 |
| -1.7947 | 93.812 |
| -2.1111 | 120.57 |
| -1.681 | 100.11 |
| -1.188 | 74.141 |
| -2.3496 | 79.337 |
| -1.2448 | 62.002 |
| -1.4008 | 50.957 |
|  | 109.04 |
| -1.6209 | 124.2 |
| -1.1969 | 83.061 |
| -1.8607 | 100.11 |
| -1.791 | 77.576 |
|  | 28.675 |
| -1.5517 | 72.891 |
| -1.9671 | 132.08 |
|  | 91.961 |
| -2.0639 | 53.013 |
| -1.8136 | 118.51 |
| -2.0652 | 116.73 |
| -1.3561 | 106.32 |
| -1.2664 | 126.07 |
|  | 65.179 |
| -2.042 | 114.4 |
| -1.9194 | 125.67 |
| -1.6522 | 47.712 |
| -1.9426 | 134.26 |
| -1.8609 | 126.12 |
| -1.7304 | 84.053 |
| -2.2966 | 69.275 |
| -1.2506 | 107.97 |
| -1.7486 | 138.98 |
| -2.1871 | 75.088 |
| -2.1729 | 80.555 |
| -2.5458 | 143 |
| -1.1672 | 164.77 |
| -2.5885 | 110.31 |
| -1.6105 | 102.07 |
| -1.8224 | 75.968 |
| -2.1984 | 154.35 |
| -1.8606 | 67.414 |
| -2.0055 | 51.445 |
| -1.7243 | 55.258 |
| -2.0949 | 120.62 |
| -2.3797 | 90.777 |
| -1.2415 | 180.78 |
| -2.1355 | 140.09 |
| -1.6822 | 131.35 |
| -2.6325 | 127.68 |
| -2.9299 | 120.9 |
| -2.8239 | 98.523 |
| -1.7243 | 67.507 |
| -1.6582 | 113.53 |
|  | 71.085 |
| -2.561 | 74.376 |
| -2.0103 | 77.899 |
|  | 93.178 |

**Supplemental Figure S3. Pie charts of the distribution of 2-hydroxyisobutyrylation proteins based on their predicted subcellular localization (A), cellular components (B), molecular functions (C), and biological processes (D).**

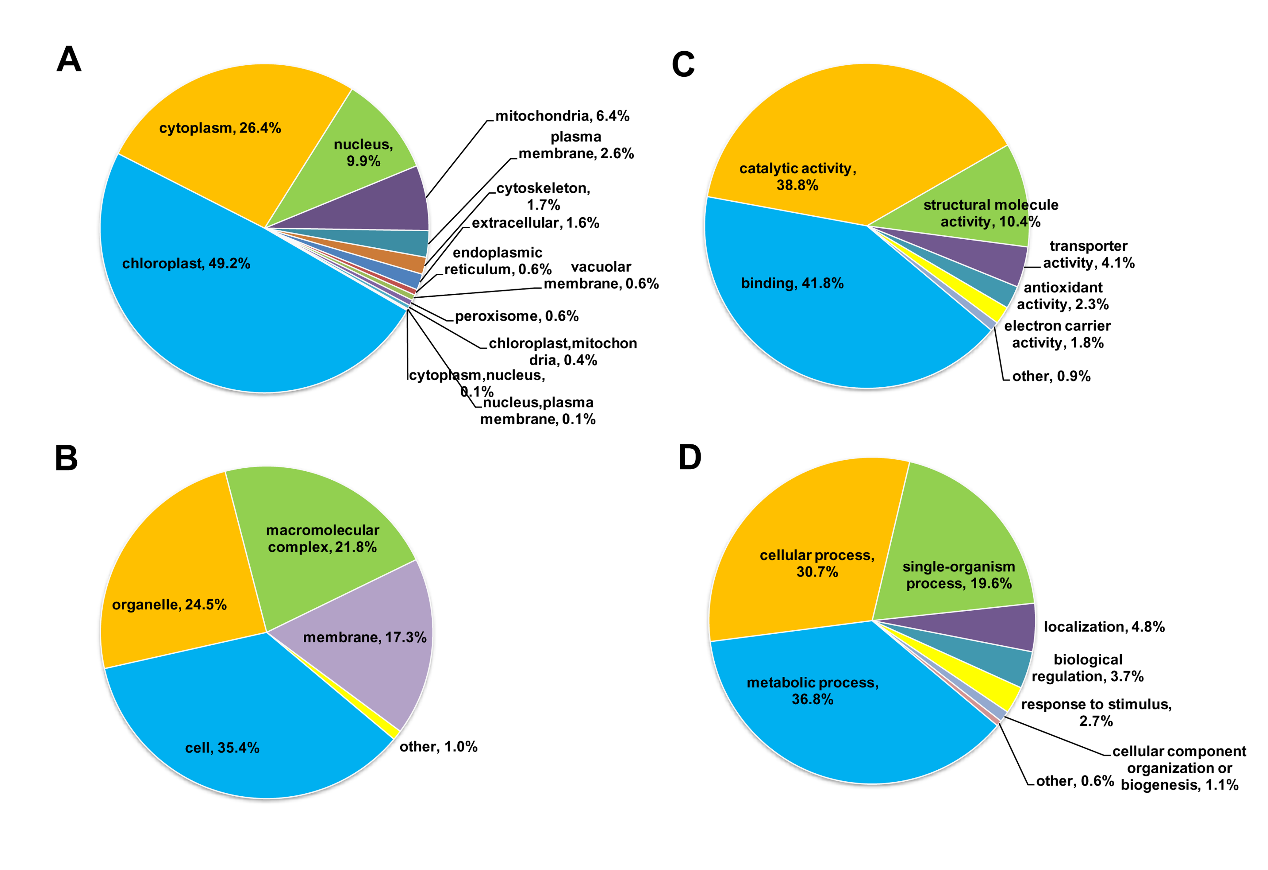

**Supplemental Figure S4. Interaction networks of the 2-hydroxyisobutyrylation proteins in wheat using String software.**

(A)	102 ribosome-related proteins were mapped the interation. (B) 11 proteins associated with proteasome were mapped the interation. (C) 10 proteins involved in metabolic were mapped the interation. (D) 435 proteins were mapped the interation.

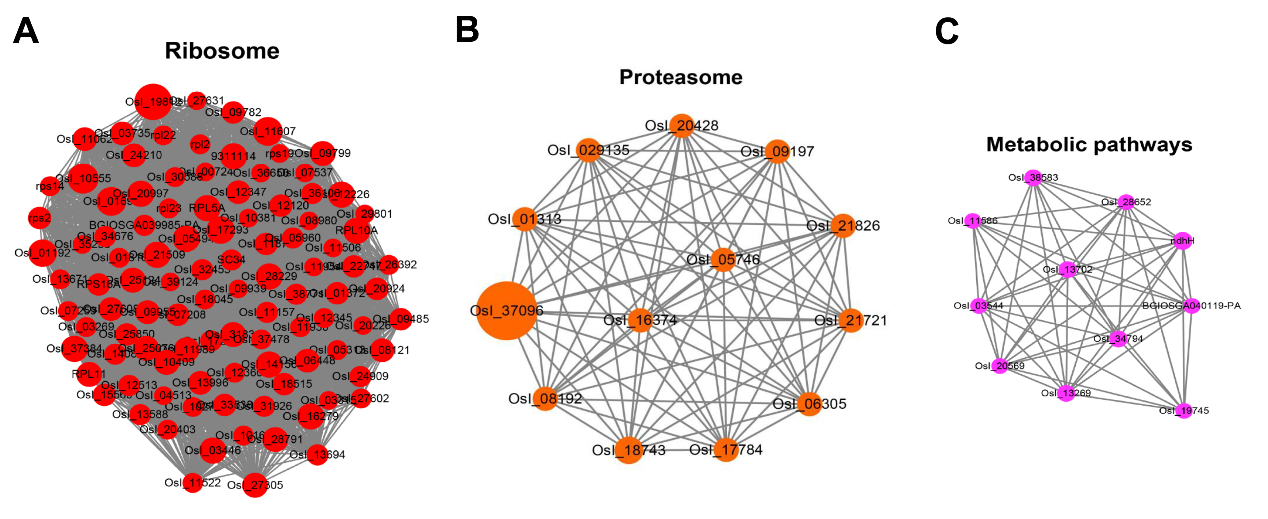

**D**

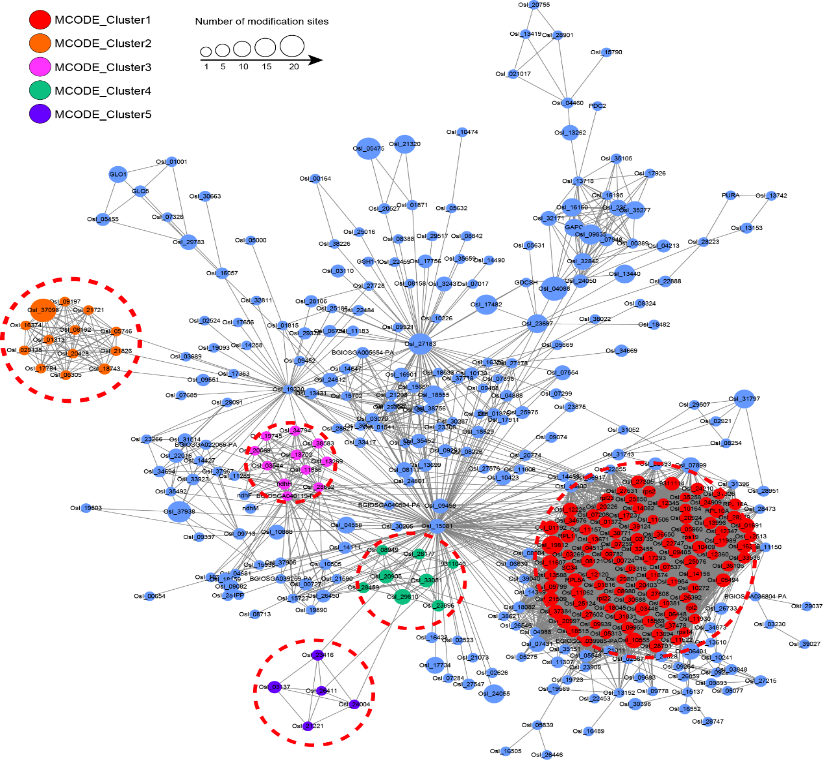

**Supplemental Figure S5.** **The tertiary structure prediction information of 14-3-3 (****L0GED8) using SWISS-MODEL.**
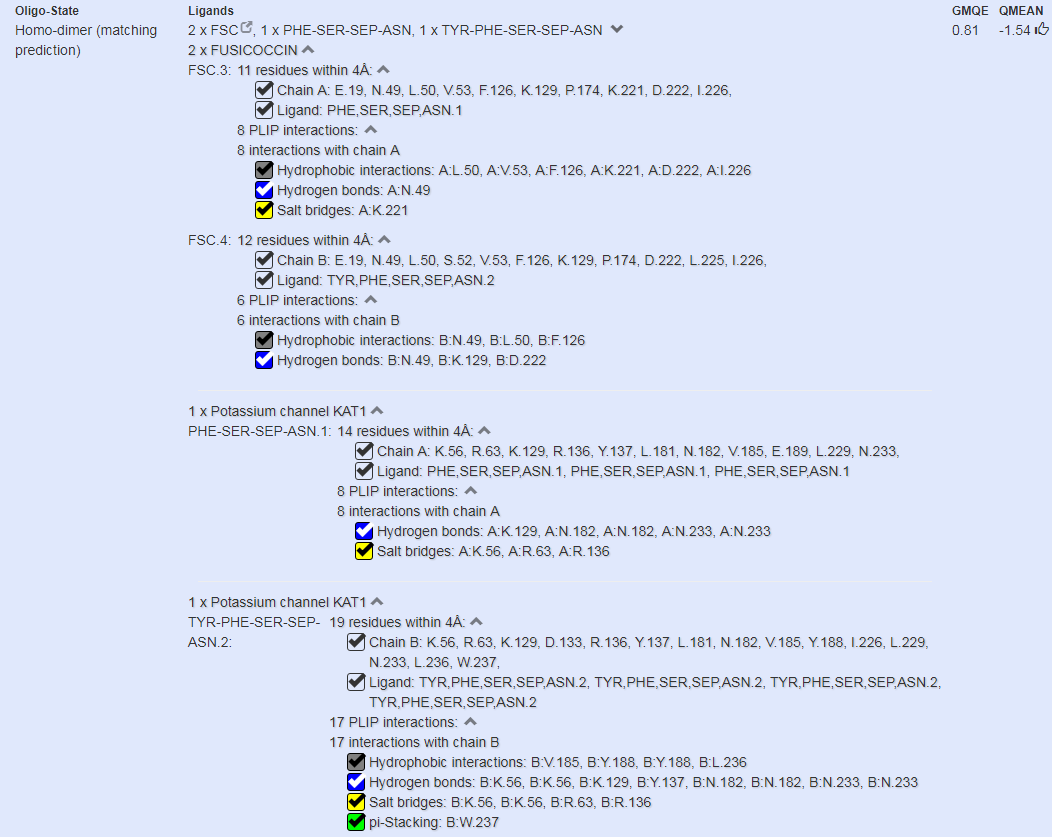

**Supplemental Figure S6. 2-hydroxyisobutyrylation proteins involved in Photosynthesis.**

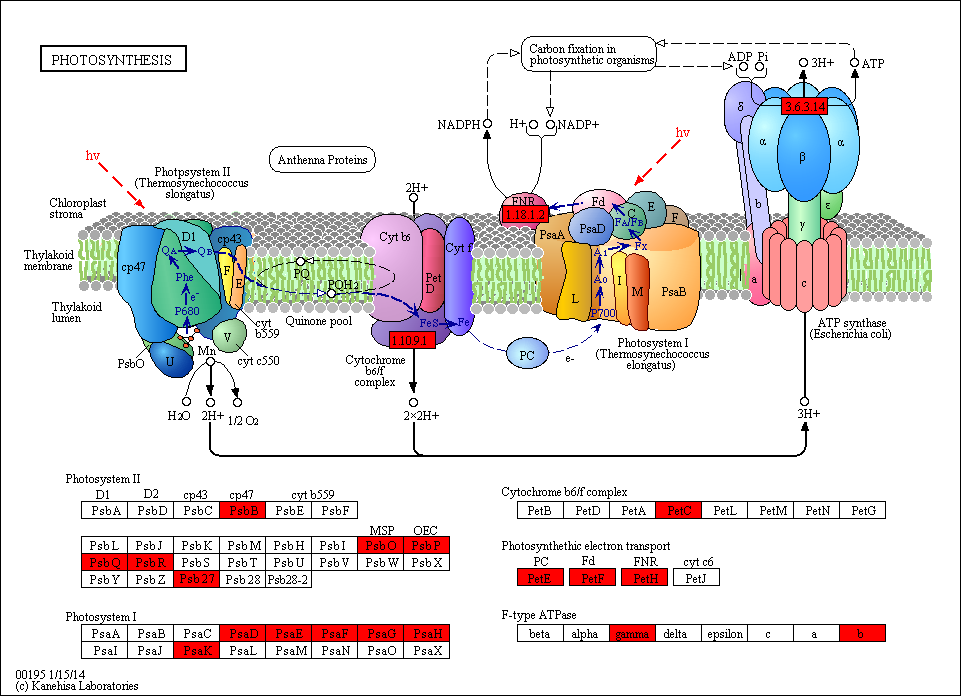

**Supplemental Figure S7. Location of Lysine,** **acetylation, succinylation, malonylation, 2-hydroxyisobutyrylation, and** **phosphorylation sites in wheat large subunit of Rubisco (RbcL, P11383).**

Kac: acetylation; Ksu: succinylation; Kma: malonylion; Khib: 2-hydroxyisobutyrylation; T34, S208, T474 were phosphorylation sites; T: Threonine, S: Serine.

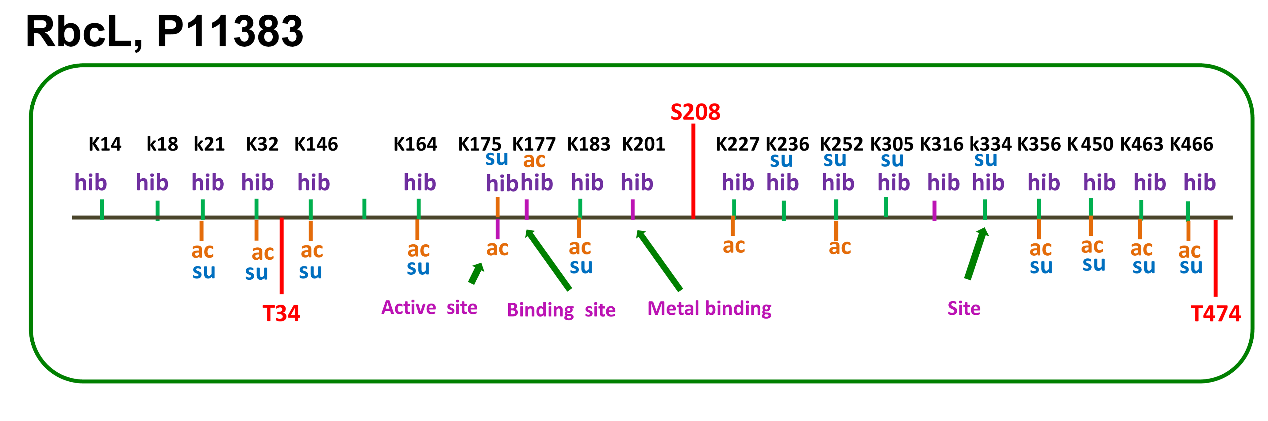

**Supplemental Figure S8. 2-hydroxyisobutyrylation proteins involved in glycolysis/gluconeogenesis.**

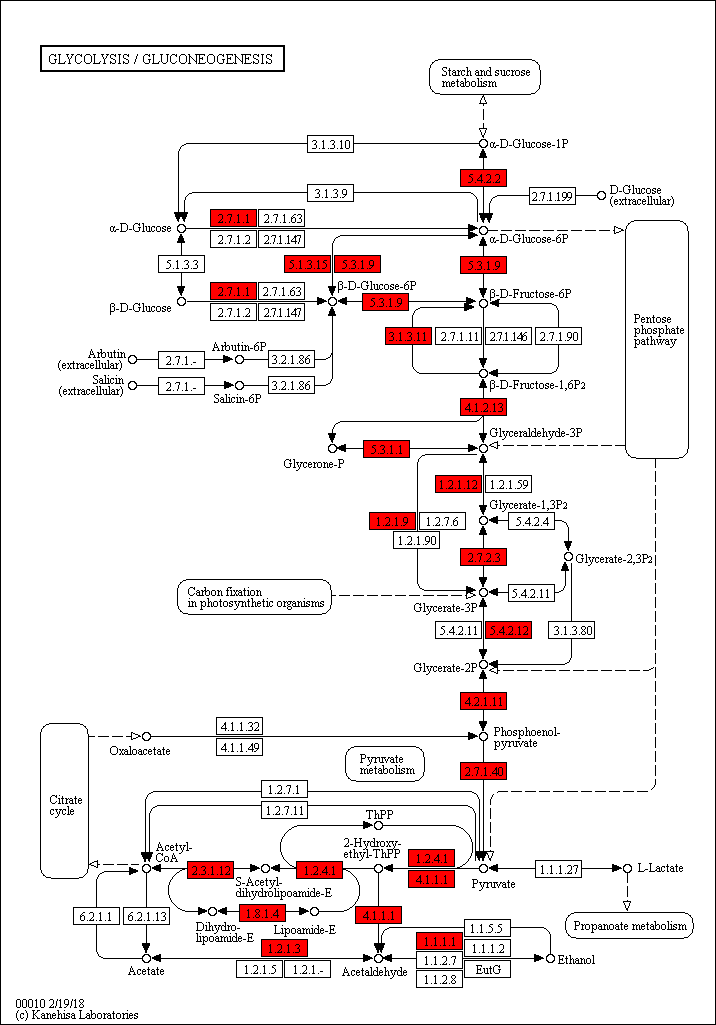

**Supplemental Figure S9. The secondary structures analysis of phosphoglycerate kinase (PGK, W5H4V7).**

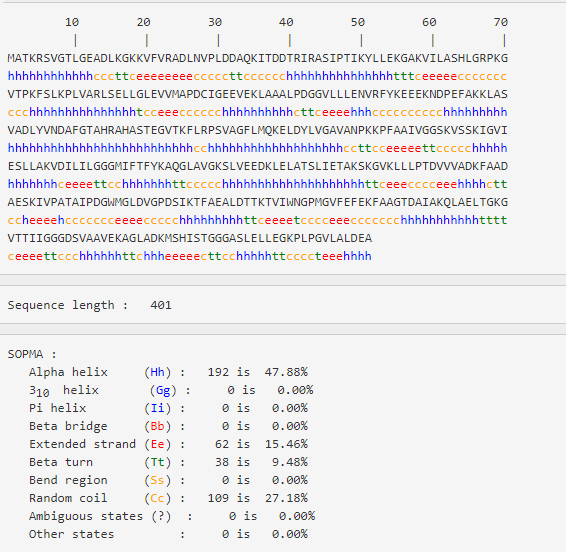

**Supplemental Figure S10. Effects of Khib on 14-3-3 with its interactive proteins.**

(A) The tertiary structure prediction of PGK constructed by SWISS-MODEL; the K206 site is located at an *α*-helix. (B) Multiple sequence alignment of 14-3-3 from different species; the conserved K56, K124, K129 site is marked by a ring. (C) [Co-IP](about:blank) of 14-3-3 (WT, K56R, K56Q, K124+129R, K124+129Q, K56+124+129R and K56+124+129Q) associated proteins. GST mAb was used. (D) LC-MS/MS analysis of 14-3-3 WT and its mutants.

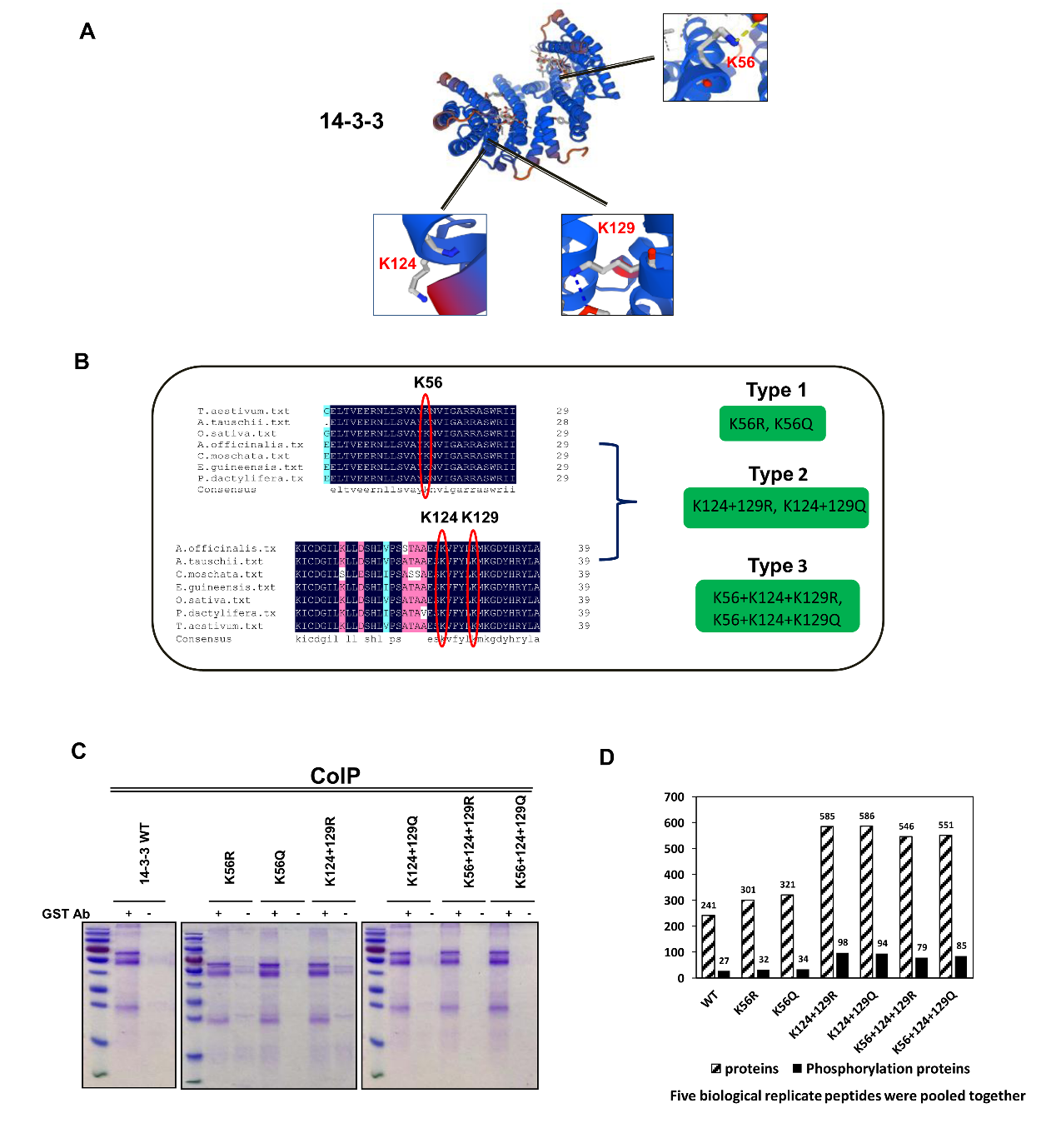
