## Supplemental Method S1 for "Global Profiling of 2-hydroxyisobutyrylome in Common Wheat"

**Supplemental Method S1: The procedure of plant protein extract, trypsin digestion,** **HPLC Fractionation, affinity enrichment, LC-MS/MS analysis and database search**

**For plant protein extract**

The sample was grinded by liquid nitrogen into cell powder and then transferred to a 5-mL centrifuge tube. After that, four volumes of lysis buffer (8 M urea, 1% Triton-100, 10 mM dithiothreitol, and 1% Protease Inhibitor Cocktail) was added to the cell powder, followed by sonication three times on ice using a high intensity ultrasonic processor (Scientz). (For PTM experiments, inhibitors were also added to the lysis buffer, e.g. 3 μM TSA and 50 mM NAM for acetylation.) The remaining debris was removed by centrifugation at 20,000 g at 4 °C for 10 min. Finally, the protein was precipitated with cold 20% TCA for 2 h at -20 °C. After centrifugation at 12,000 g 4 °C for 10 min, the supernatant was discarded. The remaining precipitate was washed with cold acetone for three times. The protein was redissolved in 8 M urea and the protein concentration was determined with BCA kit according to the manufacturer’s instructions.

样品从-80 °C取出，称取适量组织样品至液氮预冷的研钵中，加液氮充分研磨至粉末。各组样品分别加入粉末4倍体积酚抽提缓冲液（含10 mM二硫苏糖醇，1%蛋白酶抑制剂，3 μM TSA，50 mM NAM），超声裂解。加入等体积的Tris平衡酚，4 °C，5500 g离心10 min，取上清并加入5倍体积的0.1 M乙酸铵/甲醇沉淀过夜，蛋白沉淀分别用甲醇和丙酮进行洗涤。最后沉淀用8 M尿素复溶，利用BCA 试剂盒进行蛋白浓度测定。

**Trypsin Digestion**

For digestion, the protein solution was reduced with 5 mM dithiothreitol for 30 min at 56 °C and alkylated with 11 mM iodoacetamide for 15 min at room temperature in darkness. The protein sample was then diluted by adding 100 mM NH4HCO3 to urea concentration less than 2M. Finally, trypsin was added at 1:50 trypsin-to-protein mass ratio for the first digestion overnight and 1:100 trypsin-to-protein mass ratio for a second 4 h-digestion.

**HPLC Fractionation**

The tryptic peptides were fractionated into fractions by high pH reverse-phase HPLC using Thermo Betasil C18 column (5 μm particles, 10 mm ID, 250 mm length). Briefly, peptides were first separated with a gradient of 8% to 32% acetonitrile (pH 9.0) over 60 min into 60 fractions. Then, the peptides were combined into fractions and dried by vacuum centrifuging.

**Affinity Enrichment**

To enrich Khib modified peptides, tryptic peptides dissolved in NETN buffer (100 mM NaCl, 1 mM EDTA, 50 mM Tris-HCl, 0.5% NP-40, pH 8.0) were incubated with pre-washed antibody beads (Lot number PTM-804, PTM Bio) at 4°C overnight with gentle shaking. Then the beads were washed four times with NETN buffer and twice with H_2_O. The bound peptides were eluted from the beads with 0.1% trifluoroacetic acid. Finally, the eluted fractions were combined and vacuum-dried. For LC-MS/MS analysis, the resulting peptides were desalted with C18 ZipTips (Millipore) according to the manufacturer’s instructions.

**LC-MS/MS Analysis**

The tryptic peptides were dissolved in 0.1% formic acid (solvent A), directly loaded onto a home-made reversed-phase analytical column (15-cm length, 75 μm i.d.). The gradient was comprised of an increase from 6% to 23% solvent B (0.1% formic acid in 98% acetonitrile) over 26 min, 23% to 35% in 8 min and climbing to 80% in 3 min then holding at 80% for the last 3 min, all at a constant flow rate of 400 nL/min on an EASY-nLC 1000 UPLC system.

The peptides were subjected to NSI source followed by tandem mass spectrometry (MS/MS) in Q Exactive^TM^ Plus (Thermo) coupled online to the UPLC. The electrospray voltage applied was 2.0 kV. The m/z scan range was 350 to 1800 for full scan, and intact peptides were detected in the Orbitrap at a resolution of 70,000. Peptides were then selected for MS/MS using NCE setting as 28 and the fragments were detected in the Orbitrap at a resolution of 17,500. A data-dependent procedure that alternated between one MS scan followed by 20 MS/MS scans with 15.0s dynamic exclusion. Automatic gain control (AGC) was set at 5E4.

**Database Search**

The resulting MS/MS data were processed using Maxquant search engine (v.1.5.2.8). Tandem mass spectra were searched against UniProt_*Triticum* database (116722 sequences, released at 2018-04) database concatenated with reverse decoy database. Trypsin/P was specified as cleavage enzyme allowing up to 4 missing cleavages. The mass tolerance for precursor ions was set as 20 ppm in First search and 5 ppm in Main search, and the mass tolerance for fragment ions was set as 0.02 Da. Carbamidomethyl on Cys was specified as fixed modification and Lys 2-hydroxyisobutyrylation, acetylation on protein N-terminal and oxidation on Met were specified as variable modifications. FDR was adjusted to < 1% and minimum score for modified peptides was set > 40.
